# Inference of complex demographic history using composite likelihood based on whole-genome genealogies

**DOI:** 10.1101/2025.10.07.680347

**Authors:** Drew DeHaas, Zhibai Jia, Leo Speidel, Xinzhu Wei

## Abstract

Accurate parametric inference on complex demographic models is a continuing challenge in population genetics. Ancestral recombination graphs (ARGs) provide richer information than simple population genetic summary statistics and can potentially improve the power and accuracy of such inference. We present *mrpast*, a tool for inferring complex demographic history from ARGs. *mrpast* uses a composite likelihood formulation based on the pairwise sample coalescence time, observable in an ARG, and the coalescence probabilities from a continuous-time Markov process. We have evaluated *mrpast*’s accuracy on a variety of models, including stepping-stone models with asymmetric migrations, changes in effective population sizes, out-of-Africa, and American admixture. We demonstrated *mrpast*’s accuracy using simulated ARGs and inferred ARGs, and its high versatility in jointly inferring all parameters in complex models, including time of demographic events (e.g., population split, admixture), effective population size (e.g., constant, exponential growth), and gene flow (e.g., admixture proportion, migration rate). Extending the three-population out-of-Africa model with asymmetric migrations, we observed significantly more migrations from East Asians to Europeans than from Europeans to East Asians. Notably, *mrpast* can reliably recover all parameters in an American admixture model, when treating non-admixed Native Americans as an unsampled (“ghost”) population. Applying this model to Mexican, Puerto Rican, and Colombian populations, we found that the reconstructed histories of Native and admixed Americans align closely with both historical records and genetic evidence. Lastly, *mrpast* provides a comprehensive pipeline to facilitate easier, more appropriate, and robust demographic inference, in which users can easily simulate, infer, and manipulate ARGs, and illustrate and test a demographic model.

## 1 Introduction

Developing and applying methods for inferring demographic history has been a central theme in population genetics. Genetic data can reveal important information about a species’ past and shed light on a species’ future (e.g., extinction risk). Inference of recent human demographic history from genetic data offers an independent line of evidence that complements archaeological findings to provide a more comprehensive view of human history. An accurate demographic model can shed light on the dynamics of deleterious and disease variants [48], the action of natural selection [9], and the transferability of polygenic scores [13].

Generally speaking, demographic inference can be model-based or model-free. Model-free methods such as principal component analysis (PCA) ([40]), pairwise sequentially Markovian coalescent (PSMC) [33] and STRUCTURE ([45]) have been used as a de facto standard in the exploration of population genetic data [32]. These methods make fewer assumptions about the data generating process (i.e., demography), and their results can often be robust. However, results from these methods could be over/miss-interpreted (e.g., STRUCTURE [32]) and they do not provide fine-grained demographic information. In contrast, model-based parametric methods make specific assumptions about the demography, yielding estimates of more informative parameters, such as the time and intensity of gene flow. Although highly interpretable, such parametric inference requires substantial domain knowledge about the study population to set up properly, and fully parametric methods are typically restricted to simpler demographic scenarios involving only a few populations and parameters [21, 22]. Currently, parametric methods that can scale to more than a dozen populations have to make simplified assumptions to remain computationally tractable. For example, EEMS and MAPS assume local similarities of demographic parameters and symmetric migrations [4, 43], and TreeMix assumes that a tree topology describing population split history can be fixed first before inferring potential admixture events [44]. These methods utilize genetic summary statistics at the haplotype level or the population allele frequency level, such as pairwise genetic distance (e.g. EEMS [43]), length of identity by descent (IBD) segments (e.g. MAPS [4]), site frequency spectra (SFS) (e.g., δaδi [22], fastsimcoal2 [14], and momi2 [28]), and allele frequency covariance (treemix [44]). However, summary statistics such as the SFS capture only low-dimensional information from genomes, and SFS-based population size estimators converge much more slowly with increasing data than many classical estimators in statistics [51]. While haplotype-based method might capture more information than the SFS, it is harder to avoid using regions influenced by linked selection, leading to biases in demographic inference [20].

More recently, there has been a paradigm shift from developing demographic inference methods based on summary statistics to ones based on the ancestral recombination graph (ARG) [15, 19, 42]. The ARG is a model that comprehensively captures the interwoven genealogical histories of individuals along the genome. Driven by increasing genomic data availability and computational power, ARG inference methods have recently seen significant advances in scalability [29, 49]. ARGs can provide significantly more granular information in the form of genealogical trees along the genome than other summary statistics. Two recent ARG-based methods by Osmond et al.[42],and Grundler et al. [19] attempted to map ancestral haplotypes to a spatio-temporal grid, the former by estimating dispersal rates using a Brownian motion model, and the latter by explicitly mapping nodes in trees to geographical locations via a parsimony approach assuming minimum migration. Another method, gLike [15], computes the full likelihood of a marginal coalescent tree to infer parameters in models with a pulse-like admixture event. It maps nodes in a coalescent tree (which have coalescence times associated with them) to the possible population memberships of the lineages that coalesce at that node and computes the marginal probability of each state that a node can have. The likelihood of the tree is then the sum of the probabilities of the root states. Interestingly, gLike could jointly estimate ten to twenty parameters in recent admixture models using as few as ten sparsely sampled trees along the genome, suggesting that coalescent trees are highly informative about demography.

Subpopulations interconnected by continuous migration are often considered realistic for natural populations. Currently, ARG-based demographic inference has not been extended to models with continuous migration, and existing parametric methods are generally limited to jointly inferring parameters for only a few populations. Developing a more scalable method that incorporates continuous migration parameters would broaden the scope of genealogy-based inference beyond estimates of dispersal rates or admixture proportions and improve the flexibility of model-based inference. Moreover, continuous space populations can be approximated by many-deme models with migration between adjacent demes (i.e., the stepping-stone model) [43], and admixture can be represented by epochs of migration [25]. Furthermore, it remains an open question whether the rich information encoded in ARGs can be leveraged to accurately infer migration rates, especially asymmetric migration, which is particularly challenging with current summary statistic methods.

Here we present *mrpast* (*m*igration *r* ate and *p*opulation size *a*cross *s*pace and *t* ime), an ARG-based demographic likelihood method that can work with complex models containing continuous migrations and admixtures. In particular, our method leverages information from whole-genome genealogical trees while maintaining scalability, making it well suited for capturing subtle signals in present-day genomes and for analyzing datasets with limited samples per population. This flexibility enables the exploration of a wide range of species and demographic scenarios.

## 2 Results

### 2.1 Overview of Method

*mrpast* uses a likelihood formulation based on coalescence times extracted from ARGs. An ARG can be viewed as a sequence of marginal coalescent trees, each of which spans a region of the genome separated by recombination. The coalescence time *T*_*coal*_(*i, j*) between a pair of haploid samples *i, j* can be extracted from such coalescent trees. For computational tractability, we discretize continuous time ([0, ∞)) into *t* non-overlapping slices, and each coalescence time falls into a time slice. The probabilities of coalescence under a given model are obtained from a continuous-time Markov chain capturing the state of a pair of lineages (similar to the rate matrix formulations from [39] and [5]). There are 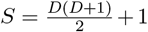 states in the Markov chain: a lineage pair can occupy any pair of the *D* demes, plus an absorbing state representing coalescence. Under a given model, the *t*-by-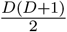 matrix **P** is defined such that **P**_(*k*,Γ(*α,β*))_ is the probability mass that two lineages starting in state 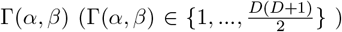 have coalesced in time slice *k* (*k* ∈ {1, …, *t*}). The matrix **C** is defined such that **C**_(*k*,Γ(*α,β*))_ is the number of observed events that a pair of lineages that started in demes *α, β* have coalesced in time slice *k*. We factorize the likelihood across pairwise coalescence events, effectively treating them as independent, and formulate the composite likelihood function

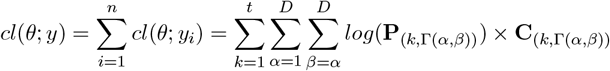

Here, *θ* represents the set of unknown parameters, which in this case, include effective population size, migration rate, growth rate, admixture proportion, and end-of-epoch time. *y* = (*y*_1_, …, *y*_*n*_) represents the observed coalescence events, where each *y*_*i*_ is a single coalescence assumed to be independent and identically distributed (i.i.d.). We use trees sampled from the ARGs to generate the observed coalescence times, and then maximize the composite likelihood *cl*(*θ*; *y*) to solve for *θ* that best fit the data.

*mrpast* is designed to work with a variety of models. As input, *mrpast* requires a model that defines the paths of migration between demes, as well as any population splits (e.g. deme *α* starting as an offshoot of deme *β*). In the model, symbolic parameters are assigned to migration rates, growth rates, admixture proportions, and effective population sizes for each deme and epoch. These parameters can be shared across epochs or can be epoch-specific. The transition times between epochs can also be parameterized. A Markov chain is constructed for each epoch of the model, which is used to compute **P**_(*k*,Γ(*α,β*))_. Migration between demes can be asymmetric if the two directions are parameterized independently. Once the parameters in a model have taken numeric values (either via inference or specified by a user), the model can be used to simulate data (Figure 1). Hence, the accuracy of resolving a specific model and the resulting uncertainty in the parameters (e.g., via parametric bootstrap) can be tested via simulation. *mrpast* supports the following three typical modes of operation:

**Figure 1:**
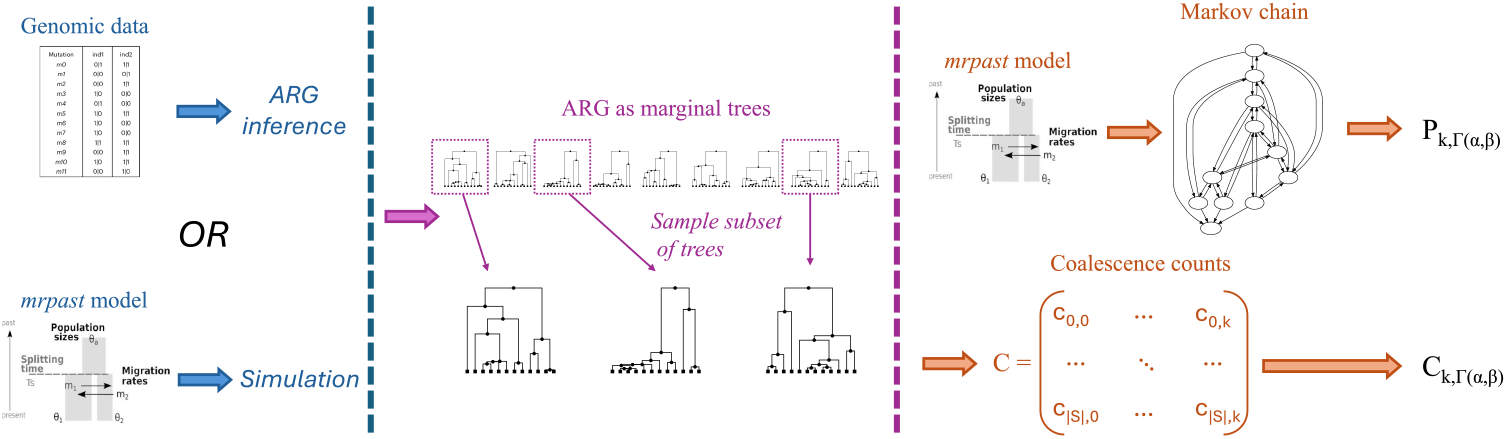
An overview of *mrpast* workflow.

A. Simulate ARGs from the model, and use coalescence times from simulated ARGs to generate **C**.
B. Simulate ARGs from the model and output the genetic data in the variant call format (VCF). Then use coalescence times from ARGs reconstructed from the genetic data to generate **C**.
C. Use empirical/real genetic data as input to ARG inference, and then use coalescence times from those ARGs to generate **C**.

Through integration with the coalescent simulator *msprime* [6] and a number of ARG inference tools [29, 49, 53], *mrpast* enables users to validate the feasibility of resolving their model of interest. If parameters in a particular model cannot be inferred with acceptable accuracy from simulated ARGs (mode *A*) or ARGs reconstructed from simulated data (*B*), then it will likely not perform well on real data (mode *C*).

### 2.2 Overview of Applications

Here, we choose four models to represent the variety of scenarios *mrpast* could be applied to: stepping-stones, changes in effective population size, Out-of-Africa Migration, and American admixture. We used a synthetic example for the stepping-stones model, two models from *stdpopsim* [3, 16, 22, 27, 50], and constructed a new model to study the history of American admixtures. Featured among these models are multiple epoch times, continuous migration, exponential population growth, admixture, and up to one unsampled “ghost” population. In the results shown, all parameters within a model are inferred jointly, although the framework also allows for fixing certain parameters to predefined values.

For each model, we used *mrpast* to call *msprime* to generate simulated ancestral recombination graphs (ARGs) in the *tskit* tree sequence format and corresponding genetic data as VCF files. Each simulated dataset has 20 100 Mbp regions, mimicking the size of the human genome. Results from inferred ARGs are based on *tsinfer* [29] and *tsdate* [53] with branch length estimated using the recently available “variational gamma” method implemented in *tsdate* [2].

#### 2.2.1 Stepping-stones Model

Stepping-stone models arrange demes in a spatial pattern, with exchanges occurring between neighboring demes, making them useful for studying populations in continuous space. Here, we created a synthetic model to test *mrpast*’s ability to study spatial models and to work with a large number of populations. We setup 20 demes in a 4-by-5 squared grid (Fig. 2a), where all 62 migration parameters are randomly sampled from a uniform distribution, hence migration between neighboring demes is asymmetric (i.e. migration from deme *i* to *j* is different from migration from deme *j* to *i*). Furthermore, each deme has a randomly sampled effective population size (20 *N*_*e*_ parameters). In this example, 20 diploid genomes per deme from simulated ARGs are used for inference with the optimization bound [0.1*x*, 10*x*] when *x* is the true parameter. We either jointly infer all 82 parameters (Fig. 2b), or infer 62 migration parameters conditional on the ground truth *N*_*e*_ (Fig. 2c).

**Figure 2:**
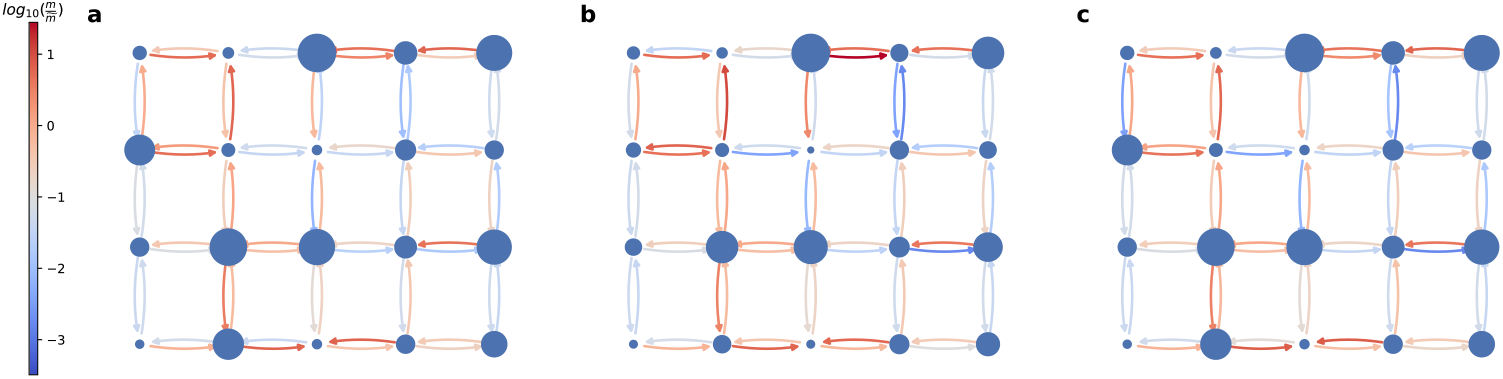
Results on stepping stone model with simulated ARGs. **a**: true values for effective population sizes *N*_*e*_ (reflected by node size) and asymmetric migration rates between populations (reflected by edge color). **b**: *mrpast* results with all *N*_*e*_ and migration rates as parameters. **c** *mrpast* results with all *N*_*e*_ values fixed, and only the migration rates as parameters.

Key factors of interest include the presence of migration barriers (low migration rates in both directions), regions with elevated migration rates (high migration rates in both directions), and strongly directional migration (e.g., north-to-south or east-to-west). We found that *mrpast* can accurately resolve many of these asymmetric migration rates and population sizes (Fig. 2a), and fixing *N*_*e*_ parameters does not noticeably improve the inference of migration rates (Fig. 2b).

#### 2.2.2 Changes in Effective Population Size

The inverse instantaneous coalescent rate (IICR) over time (often interpreted as changes in effective population size) can provide important information about the evolutionary history and long-term viability of the study population; IICRs of multiple subpopulations could be compared against each other to understand demographic history. IICR can be estimated from DNA sequences using programs such as PSMC (pairwise sequentially Markovian coalescent) [33], or from pairwise coalescence times directly if ARGs are available [49]. Although IICR can be directly interpreted, a parametric model can still offer better interpretability.

Here, we use the *Africa_1T12* model from *stdpopsim* as an example, which was originally simplified from the Tennessen et al. model [3, 50] to model the African population in isolation.

This is a single population three-epoch model, with exponential growth in the most recent epoch, and constant effective population sizes in the other two epochs (Fig. 3a). We generated 50 replicate simulation datasets, each consisting of 100 diploid genomes. Using simulated ARGs, we found that *mrpast* produces nearly unbiased estimates (Fig. 3bc), with relative errors for the median parameter values (over all replicates) within 17%, except for the most recent (largest) *N*_*e*_. We then performed ARG inference using the first simulated dataset and generated 100 bootstrapped datasets to assess the uncertainty. The estimates from inferred ARGs (Fig. 3de) are mostly comparable to those from simulated ARGs (Fig. 3bc), and the patterns of variation are consistent as well.

**Figure 3:**
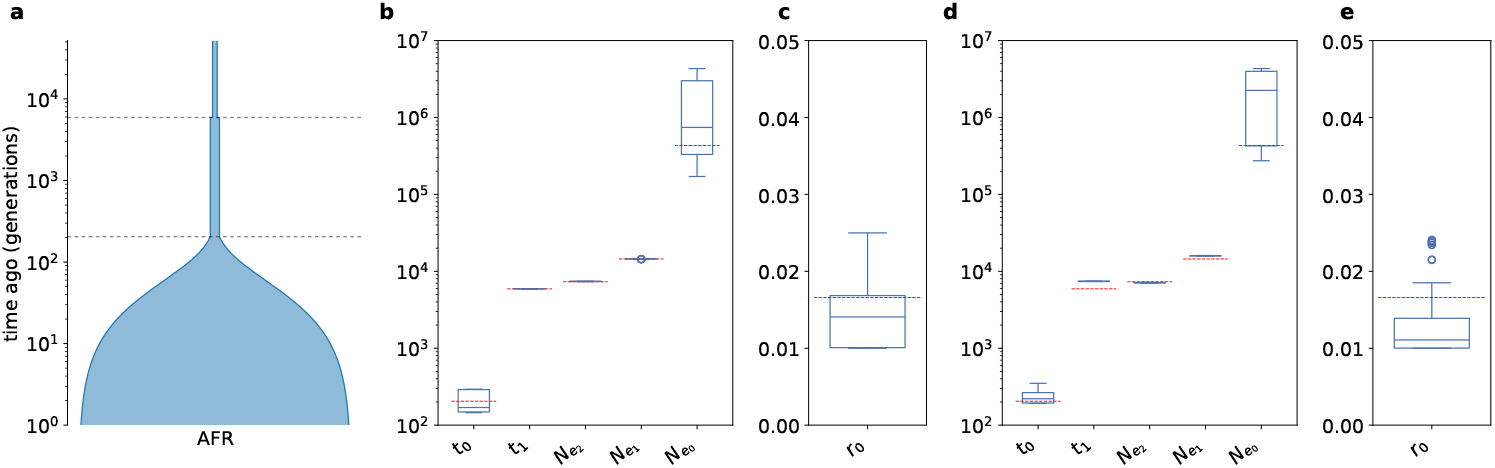
Results on population growth model. Diagram of the model structure in panel **a**. Panel **b** shows estimated parameter values on simulated ARGs, for time and effective population size. Panel **c** shows growth rate on simulated ARGs. The variation shown is from 50 different simulations. Panels **d** and **e** show results on inferred ARGs for a single simulated replicate; the variation shown is from 100 bootstrap samples taken from the ARGs. For panels **b**-**e** the red dashed lines are the true simulated parameter value. Boxes span the interquartile range (IQR), whiskers are 1.5 *× IQR*, outliers shown as circles.

#### 2.2.3 Population Divergence with Migration

The isolation with migration (IM) model offers a valuable framework for studying population divergence. While the classic IM model describes a scenario in which a single population splits into two with continuous gene flow between them, it can be extended to accommodate more complex demographic histories, such as variable population sizes. Here, we use the *OutO-fAfrica_3G09* (OOA3G09) model from *stdpopsim* [3], to test the ability of *mrpast* to resolve population divergence time, migration, and growth. Originally proposed by Gutenkunst et al. [22], this model uses 14 parameters to describe the history of three populations (YRI: African, CEU: European, CHB: Chinese). This is a four epoch model with three epoch transition times, two corresponding to population splits (YRI-CEU, and CEU-CHB), and one reflecting a change in the YRI effective population size. We found that all 14 parameters in this model could be jointly estimated with high accuracy using both simulated ARGs (Fig. 4b, Pearson *r >* 0.99), and ARGs reconstructed by *tsinfer* (Fig. 4c, Pearson *r >* 0.99). Accuracy only reduces minimally when changing the sample size from 50 genomes per population (Fig. 4bc) to 10 genomes per population (Fig. S1ab).

**Figure 4:**
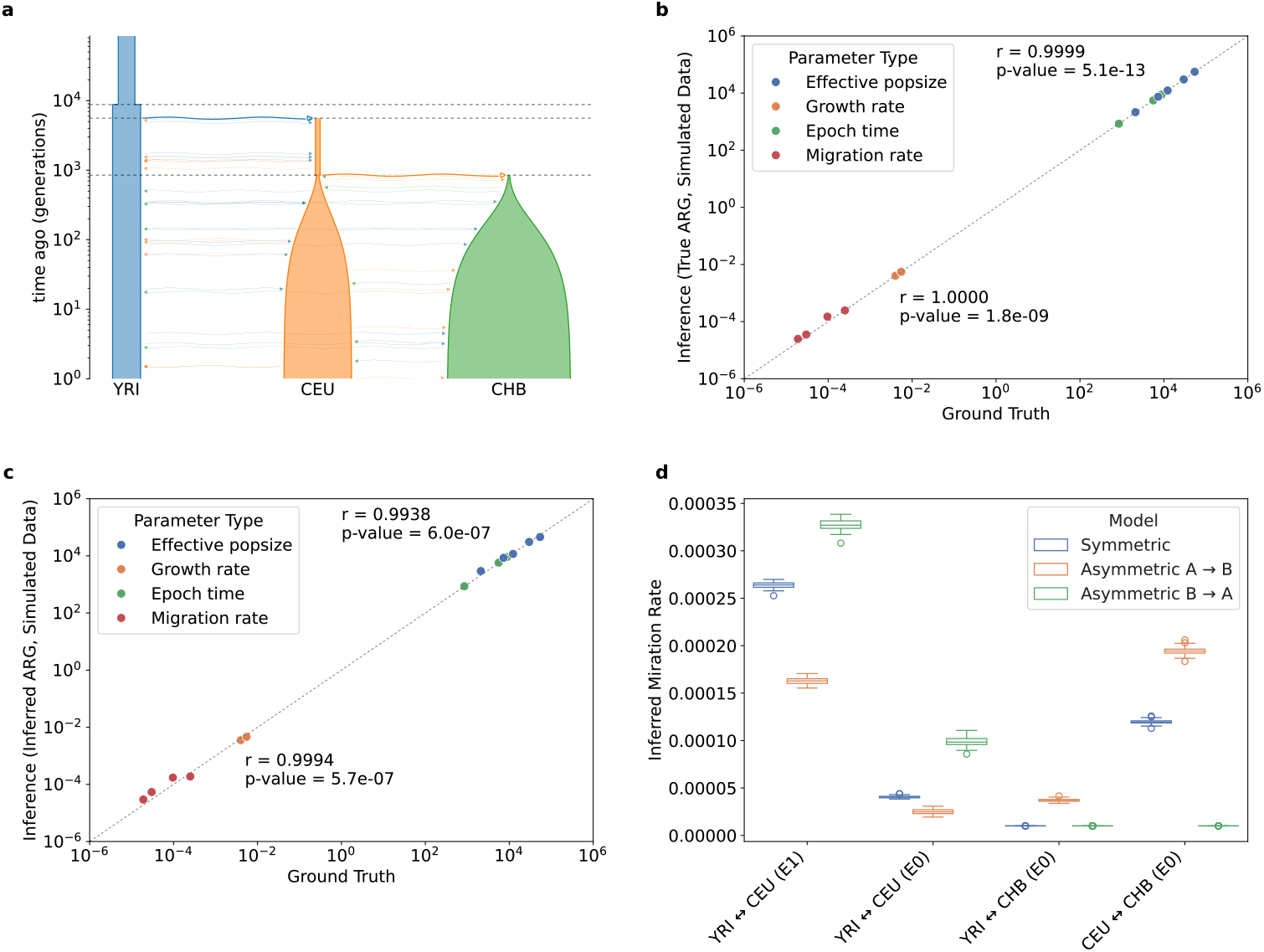
Results on three-population out-of-Africa model (stdpopsim OutOfAfrica 3G09). Diagram of the model structure in panel **a**. Panels **b** (Simulated ARGs) and **c** (Inferred ARGs) show the true (simulated) values against the inferred values on a log-log scale. The Pearson’s *r* and associated p-value are shown separately for the time and population size parameter (upper) and the rate parameters (lower). **d**: Migration rates inferred from 1,000 Genomes Project data using the symmetric migration model (OutOfAfrica 3G09) and a model modified to use asymmetric migration (OOA3G09_AM). X-axis labels describe groups of migration rate parameters *A* ↔ *B* (i.e., *A* is the first population named in the label). The asymmetric backward-in-time migration directions are described in the legend, with *A* → *B* representing the backward-in-time migration rate from A to B. E0 and E1 represent the two most recent time epochs. Boxes span the interquartile range (IQR), whiskers are 1.5 *× IQR*, outliers shown as circles.

Because the original OOA3G09 assumes symmetric migration, we further created an extension of it which assumes asymmetric migration (OOA3G09_AM) (see simulation results Fig. S2). We apply *mrpast* to ARGs inferred from the 1000 Genomes Project (KPG) to estimate parameters in OOA3G09 and (OOA3G09_AM), using 35 unrelated samples from each population, and bootstrap coalescent trees to estimate the confidence intervals. In this part, we fix the epoch time based on OOA3G09 in *mrpast* inference so that the migration parameter estimates can be compared directly. We found that three of the four symmetric migration rates falls between the corresponding pair of asymmetric migration rates. Migration rates were highest right after the out-of-Africa event, and the backward-in-time migration rate from CEU to YRI is higher (CEU→YRI(E0) Fig. 4d) than the reverse direction. The backward-in-time CEU to YRI migration rate can be interpreted as the probability that a CEU lineage has a parent/ancestor in YRI population in the previous generation. Possibly, after the out-of-Africa bottleneck, even a modest number of African migrants could have resulted in a high CEU→YRI migration rate. Under the OOA3G09_AM model, the backward-in-time migration from CEU to CHB (CEU→CHB) is much higher than the reverse in the most recent epoch. A number of historical factors could have contributed to this asymmetric migration (see Discussion and Supplemental Fig. S8).

We also replaced the CEU samples with the same number of FIN samples, and reran the OOA3G09_AM model inference (Supplementary Table S4). We observed a lower YRI to FIN migration rate (backwards in time) during the most recent 800 generations, as well as a larger FIN to CHB migration rate in the same time frame. Aside from the FIN effective population size and growth rate in the most recent epoch being smaller, the model results are otherwise consistent with the dataset utilizing CEU samples.

#### 2.2.4 American Admixture Model

Our American Admixture model describes the genetic ancestry of present-day admixed American populations as the result of recent contributions from three continental source populations: European, African, and Native American. Due to centuries of colonization and contact with Europeans and Africans, very few indigenous American groups remain isolated or scarcely admixed [17]. Although surrogate sources may be used (e.g. using East Asian as a substitute [7]), such proxies nevertheless differ from the true ancestral population and thus bias the inference. To this end, we are interested in reconstructing the demographic history of Native American (NAT) and admixed American (ADMIX) populations while treating NAT as an unsampled (“ghost”) population. To investigate this, we extended OOA3G09 (Fig. 4a) with an additional split event between Asian and Native American populations and added an admixed American population using European, Native American, and African sources (Fig. 5a). For simplicity, we assume constant effective population sizes for both Native American and admixed American populations. Compared to OOA3G09, our American admixture model introduces two additional epoch-time parameters (six epochs total), two admixture-proportion parameters, and two additional effective population sizes.

**Figure 5:**
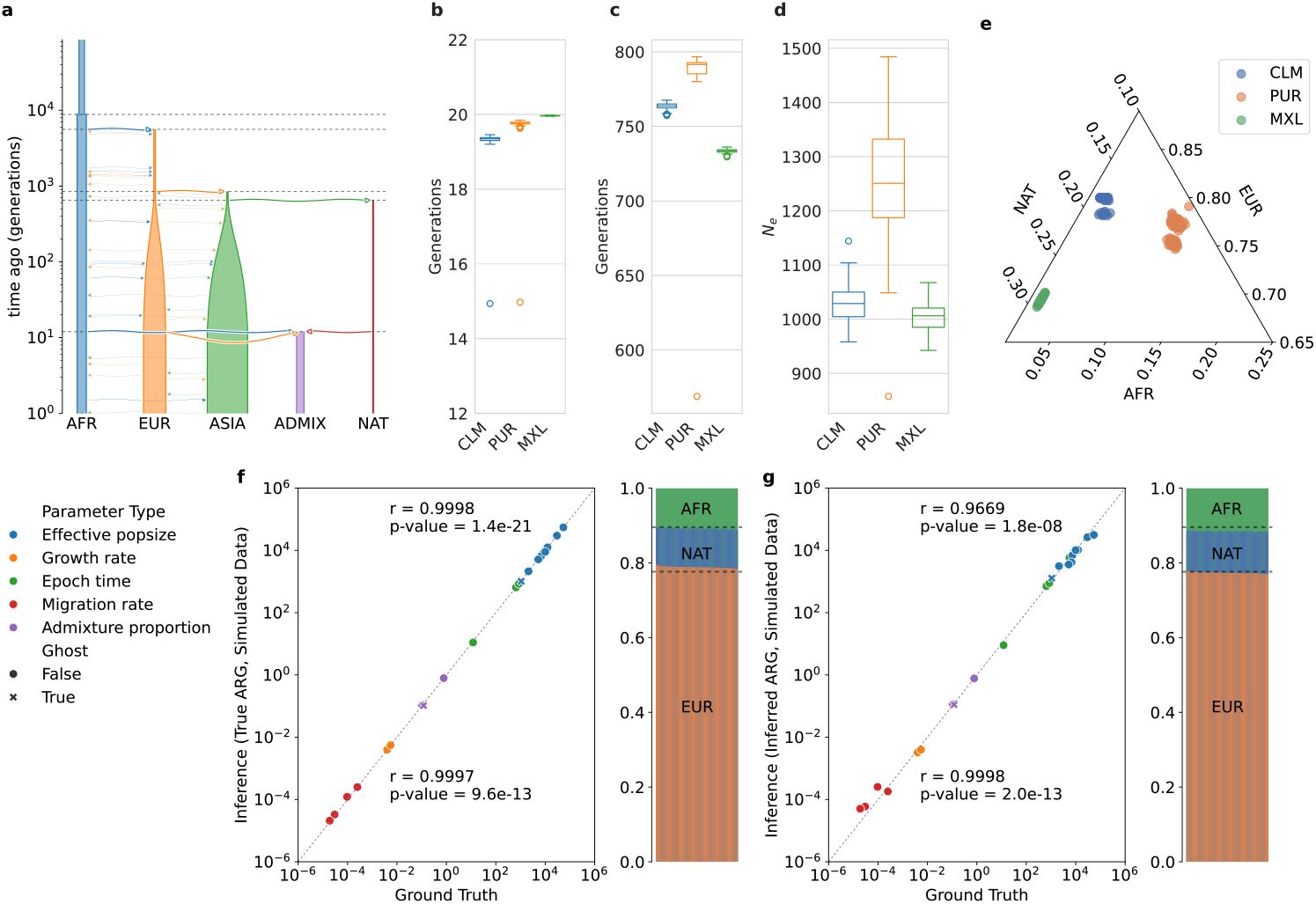
American admixture inference. **a**: Diagram of the model structure. **b**: Inferred time (generations ago) of admixture. **c**: Inferred time (generations ago) of NAT split from ASIA. **d**: Inferred effective population size for NAT. **e**: Inferred proportions of admixture for the CLM, PUR, and MXL populations. **f** : The true (simulated) values against the inferred values on a log-log scale (left), and the true vs. inferred admixture proportions (right), using simulated ARGs. The dashed lines represent the truth. **g**: Same as panel **f**, except using inferred ARGs from simulated data. 100 bootstrap samples were used in panels **b**-**g**. For panels **b, c, d**: boxes span the interquartile range (IQR), whiskers are 1.5 *× IQR*, outliers shown as circles.

Here, we use *mrpast* to jointly infer all 20 parameters including the admixture time and proportions without relying on an isolated Native American reference population – NAT is in the demographic model (Fig. 5) and used in constructing the Markov chain (Fig. 1), but only AFR, EUR, ASIA, and ADMIX samples are used for ARG inference and model fitting. Yoruban (YRI), Central European (CEU), and Han Chinese (CHB) samples from the 1000 Genomes Project are used to represent African (AFR), European (EUR), and Asian (ASIA) populations, respectively, while Mexican Americans (MXL), Puerto Ricans (PUR), and Colombians (CLM) serve as the ADMIX (Fig. 5). We fit the model to one ADMIX population at a time, using 35 unrelated samples from each population.

First, we verified that the parameters for AFR, EUR, and ASIA shared with OOA3G09 produce estimates consistent with those obtained under OOA3G09 (Supplementary Fig. S6). We estimate admixture times of 19–20 generations for CLM, PUR, and MXL (Fig. 5b). The split between East Asians and Native Americans is dated to 730–790 generations ago (Fig. 5c). In addition, under the constant effective population size model (Fig. 5a), the Native American effective population size is estimated to be only 1,000–1,300 (Fig. 5d). Finally, we find that European ancestry is higher in CLM and PUR, whereas MXL has higher Native American ancestry than PUR and CLM. These estimated admixture proportions are in strong agreement with previous studies that also used the 1000 Genomes Project data (Table 1).

**Table 1:** Comparisons of *mrpast* Admixture Proportions with Previous.

| Data | *mrpast* |  |  | Median Ancestry Prop. [52] |  |  | ADMIXTURE Result [11] |  |  |
| --- | --- | --- | --- | --- | --- | --- | --- | --- | --- |
|  | NAT | EUR | AFR | NAT | EUR | AFR | NAT | EUR | AFR |
| CLM | 0.18 | 0.80 | 0.02 | 0.25 | 0.65 | 0.045 | 0.18 | 0.75 | 0.07 |
| PUR | 0.13 | 0.77 | 0.10 | 0.14 | 0.73 | 0.1 | NA | NA | NA |
| MXL | 0.29 | 0.69 | 0.02 | 0.47 | 0.46 | 0.038 | NA | NA | NA |

As a verification, we generated simulated data under the American Admixture model and then treated NAT as a ghost population in the inference. When bootstrapping the coalescent trees, we observed a slight overestimation of EUR ancestry using simulated ARG (Fig. 5f), and a slight overestimation of AFR ancestry using ARGs inferred by *tsinfer* (Fig. 5g). But overall, all parameters including those related to ADMIX and NAT are estimated accurately (*r >* 0.99 using simulated ARG, *r >* 0.96 using inferred ARG, see Fig. 5fg).

## 3 Discussion

In summary, we present *mrpast*, an ARG-based demographic inference method that uses whole-genome genealogical trees to reconstruct demographic histories. We demonstrate its accuracy on a range of simple to complex simulated models, as well as its ability to offer new insights from the 1000 Genomes Project data. Our method works well across a wide range of common scenarios, while being able to scale to as many as 20 distinct populations.

Extending the OOA3G09 model to allow asymmetric migration (OOA3G09_AM), we found that migration rates were highest immediately after the out-of-Africa event. In the past 848 generations, we inferred greater forward-in-time east-to-west continuous migration than in the opposite direction. This discrepancy between the eastward and westward migration signals is expected based on historical / genetic evidence of Eurasian population contact and movement. Prominently, from the second century BCE through the mid-15th century, the Silk Road facilitated the movement of both goods and people across Eurasia, creating opportunities for genetic exchange, especially in the east-to-west direction. More recently, the Mongol expansions between the 13th and 14th centuries led to substantial population movements into eastern and central Europe. When we split the most recent epoch in OOA3G09 at 100 generations ago, we found that the asymmetry in migration between CEU and CHB spans both epochs (0-848 generations ago), but is more pronounced between 100 and 848 generations ago. A possible explanation for increased migration between 100 and 848 generations ago is the expansion of Central Asian groups such as Anatolian Neolithic farmers into Europe around 7,000 BCE during the spread of agriculture[41]. When using FIN instead of CEU in OOA3G09_AM, we inferred a larger FIN→CHB migration rate (backward-in-time) than CEU→CHB (Table S4), consistent with the expectation that Finns carry higher levels of East Asian ancestry [31].

Our American admixture model (Fig. 5a) captures key features of Native American and admixed American population history. Because OOA3G09 (Fig. 4) is a nested model within the American admixture model, the consistency of parameter estimates for AFR, EUR, and ASIA with those from OOA3G09 provides confidence that the established demographic relationships among AFR, EUR, and ASIA are robust to the added model complexity. Building nested models offers a natural framework for expanding and improving model-based demographic inference. Future work could incorporate automated model space exploration through iterative model updates, similar to those used in GADMA [38] and TreeMix [44].

Under the American admixture model (Fig. 5ab), the inferred admixture times of 19–20 generations (475-600 years ago assuming 25-30 years per generation) for CLM, PUR, and MXL agree well with historical records of the first European contact with Native Americans in 1492 CE, suggesting that our method is sensitive to relatively recent admixture events even when some true ancestral sources (e.g., Native American populations) are not sampled. In comparison, gLike reported a similar but older admixture time for Latinos at 25 generations ago[15]. This modest discrepancy in estimated admixture time could be due to using different datasets, demographic models, approaches, and inference parameters. In addition to admixture time, our estimated admixture proportions also align with previous estimates of global ancestry (Table 1). Lastly, we estimated that East Asians and Native Americans split 730–790 generations ago (18250-23700 years ago, assuming 25-30 years per generation; note that this is not the same as peopling of the Americas), which is broadly consistent with the previous estimate that Native Americans diverged from Siberians and East Asians approximately 23,000 years ago [36]. Taken together, our demographic inference results are consistent with historical and genetic expectations.

More broadly, this example demonstrates that *mrpast* can perform model-based joint inference of admixture time and proportions without fully relying on non-admixed source populations, or specialized frameworks that infer one type of parameters at a time (e.g., admixture proportion, or population split time). This is possible when there are exchanges between the unsampled population and some sampled populations. By estimating parameters of missing sources based on available samples, *mrpast* can provide insights into the histories of unsampled or extinct populations, based on a small number of samples from each sampled population (e.g., 35 sampled genomes per population for the American admixture model). This ability to infer information about ghost populations varies depending on the model; in supplemental Section 1 we describe a Neanderthal admixture model, where the Neanderthal effect is too weak to be reliably detected when treating Neanderthals as a ghost population.

The scalability of *mrpast*’s inference is independent of the number of samples per population. In simulations, we generally obtain accurate results with as few as 5–10 samples per population, with only modest improvements in accuracy when larger sample sizes are used (Supplemental Fig. S1). Thus, unlike methods based on allele frequencies [22, 44] or IBD segments [7], *mrpast* does not require large sample sizes, thus it remains suitable for studying natural non-human populations with limited sample availability. Nevertheless, additional samples can still be valuable for resolving more complex demographic models, especially for recent demographic history and lower migration rates. Access to larger datasets in future studies could enable the inference of finer-scale features. For example, our American admixture model assumes a constant-*N*_*e*_ in Native Americans (Fig. 5a), and current estimates place the effective population size of Native Americans at only 1,000–1,300 under this model. Having access to more admixed samples will likely enable the inference of a variable *N*_*e*_ model, which can provide a more realistic view and precise quantification of the timing and severity of historical bottlenecks in Native Americans.

We showed that *mrpast* supports a wide range of modeling frameworks, enabling users to explore diverse demographic scenarios within a single tool. Moreover, *mrpast* software can accommodate hybrid models that combine features of different frameworks, such as stepping-stone and hierarchical structures. Such versatility opens the door to systematically comparing alternative demographic histories, identifying the models that best explain observed genetic patterns, and extending inference to previously underexplored or intermediate forms of population structure. In doing so, *mrpast* provides a unified platform for developing and testing increasingly realistic demographic models.

The results from *mrpast* relies on accurately inferred ARGs. It directly utilizes the pairwise coalescence times, which is commonly used to evaluate ARG inference methods[55]. In our tests, *tsinfer* [29] combined with *tsdate* [53] performs reliably and integrates well with our framework. While powerful, they are not without limitations. Biases in ARG inference can propagate into downstream demographic analyses, resulting in a gap between *mrpast*’s accuracy using simulated ARGs and its accuracy using inferred ARGs. *mrpast* is designed to be compatible with any ARGs in *tskit* tree sequence format [54], making it readily applicable to ARGs inferred by future methods. As ARG inference methods continue to improve, their biases are expected to reduce, which should improve *mrpast*’s accuracy with inferred ARGs.

While most models in this study required only minutes to optimize (e.g., the OOA3 model took 4.3 minutes of compute time to solve all 40 solver replicates),the stepping-stone model (20 demes, 82 parameters), took three days. The most computationally intensive step in *mrpast* is to calculate the transition probabilities for a given model, for which we have implemented the highly accurate scaling-and-squaring method for computing matrix exponentials from [35]. The complexity of this method depends on the number of states – growing with 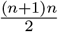 for *n* demes – as well as the number of model parameters. In our experience, reliable numerical optimization of the composite likelihood using true ARGs requires highly accurate transition probabilities. If a method with an accuracy comparable to scaling and squaring [35], but better runtime performance, becomes available in the future, it would improve *mrpast*’s scalability. Another important hyperparameter is the number of time slices used to discretize coalescence times. Once the model is specified, runtime further scales with the number of time slices and number of solver replicates. In practice, we found that models with many epochs can achieve higher accuracy when using a larger number of time slices (e.g., 100–200 under the default discretization scheme), but otherwise using 25-50 time slices is usually sufficient. The number of time slices used can be found in Table. S1. In practice, the number of time slices and solver replicates can be evaluated with simulated data to help achieve more desirable accuracy.

We have presented some comparisons of simulated vs. inferred ARGs in Figures 4 and 5 which show that results from inferred ARGs are slightly less accurate on those models. In the future it would be valuable to explore a range of models on different ARG inference methods, to explore biases that may be introduced by ARG inference. Such biases may become more obvious by looking at confidence interval coverage.

Some of the popular demographic inference methods have undergone over a decade of continuous refinement. The SFS–based method ∂*a*∂*i*, is a good example of this. ∂*a*∂*i* was first published in 2009 with support for up to three populations [22], and since then, it extended its methodology [46], improved its runtime [21], and introduced a command-line interface (CLI) [24]. Inspired by these perpetual improvements, we designed *mrpast* to be comprehensive and easy to use, supporting a variety of demographic models, methods for parameter confidence interval computation, model selection, and integration with the standard Demes format ([18]). We are optimistic that the rapid improvement to the accuracy and scalability of ARG inference will further expand the range of demographic scenarios to which techniques like *mrpast* can be applied.

## 4 Methods

### 4.1 Composite Likelihood

An ARG encapsulates the order of coalescence events among a set of haploid samples as well as the amount of time between each such event. A full likelihood function for an ARG, given a demographic model, is computationally intractable to work with. Instead, we assume independence between each observed pairwise coalescence and each sampled coalescent tree to formulate a composite likelihood function based on a subset of the information from the ARG. In practice, we utilize sparsely sampled trees along the genome that are at least 125 kilo-base-pairs (KBP) apart to make them relatively independent from each other.

Given a demographic model that has *D* demes and *E* epochs, each epoch has its own migration rates between those demes, as well as coalescent rates, population growth rates. The time of transitions between two epochs are also parameterized. Populations can be created in a more recent epoch as an admixture of populations from an older epoch. The following equations are presented as they would be solved for a single discretized time slice, and a single epoch. For details on how they are solved across all time slices and epochs, see Supplemental section 3.

Let the model parameters be **M**, a *D × D* migration rate matrix, **Λ**, a *D*-length coalescent rate vector. **M**_(*α,β*)_ is the migration rate from deme *α* to *β* (backward in time). Handling of growth rate parameters is discussed separately in the Supplemental Section 3.3.

Given a pair of lineages *i, j*, their locations in the *D* demes can be specified by one of 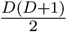 states (all unordered combinations of deme pairs). We define a continuous-time Markov chain with 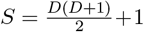 states that represent both location and coalescent state of a pair of lineages. The state 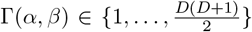 represents one of the disjoint lineages *i, j* being located in deme *α* and the other disjoint lineage being located in deme *β*, where Γ(*α, β*) = Γ(*β, α*) is unordered. Γ_*coal*_ = *S* is the absorbing state representing coalescence. Coalescence between lineages *i, j* can only occur when two lineages are in the same deme (i.e., *α* = *β*).

The transitions Γ(*α, β*) → Γ(*γ, β*) and Γ(*α, β*) → Γ(*α, γ*) are defined for pairs *α, γ* (*β, γ*, respectively) only if the input model allows for migration between the corresponding demes (e.g., **M**(**e**)_(*α,γ*)_≠ 0). We construct the infinitesimal rate matrix (“*Q*-matrix” [37]) **Q**, of size *S × S* . The transition Γ(*α, β*) → Γ(*α, γ*) (*α*≠ *β*; *β*≠ *γ*) has rate **M**_(*β,γ*)_. Γ(*α, α*) can transition to the coalescence state Γ_*coal*_ at rate **Λ**_*α*_ or transition to Γ(*α, β*) (*α*≠ *β*) at rate 2 *×* **M**_(*α,β*)_ (because *either* lineage *i* or *j* can move to *β*). We then compute the probabilities of being in each state at the end of the time slice (or epoch) via **P**_**Q**_ = *e*^**Q***t*^ (where *t* is the time elapsed since the beginning of the epoch). **P**_**Q**_ has the following structure:

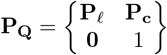

where **P**_*𝓁*_ is a (*S* − 1) *×* (*S* − 1) matrix and **P**_**c**_ is a (*S* − 1) *×* 1 matrix. **P**_*𝓁*_ represents the probability of lineage *i, j* being at a particular location at time *t*, relative to the locations at the start of the current epoch, and **P**_**c**_ the cumulative probability of having coalesced in that time. See Supplementary Section 3 for algorithms and more details, including admixture and growth rate handling.

The parameter vector ***θ*** represents the unique non-zero parameters from **M** and **Λ**, as well as growth rates, admixture proportions and epoch times. *mrpast* uses the Subplex-based ([47]) solver from NLOpt ([26]) to find 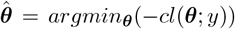. We choose the highest composite likelihood solution on multiple solver replicas with random initialization of ***θ*** to ensure the global optimum. By default, *mrpast* uses 10 solver replicates per epoch, e.g. a model with 2 epochs will use 20 solver replicates.

### 4.2 Simulations

*mrpast* integrates with *msprime* for simulation directly from the input *mrpast* model. We used 20 replicates (“chromosomes”) of length 100 mega-base-pair (MBP) each, using GRCh38 recombination rate maps from [12]. We also make use of the Demes format [18] to export models from stdpopsim and import them into *mrpast*, and many of our model topology figures were generated via DemesDraw [1].

### 4.3 ARG Processing

#### 4.3.1 ARG Inference

We used tsinfer v0.4.1 [30] with tsdate v0.2.1 [53] for all ARG inference results shown in the main text. For both the out-of-Africa 3-deme model and the American admixture model we used a mutation rate of 2.35 *×* 10^−8^ per base pair per generation, as was specified for OOA3G09 in *stdpopsim* [3].

#### 4.3.2 Sampling

We process ARG inputs in the *tskit* tree-sequence format ([29]). To account for the fact that adjacent trees are highly correlated with each other, we sample a marginal tree every *L*_*bp*_ base pairs. We used *L*_*bp*_ = 125, 000 throughout our experiments. We further restrict our sampling to regions that have a recombination rate of ≤ 1 *×* 10^−9^, with the expectation that a lower recombination rate makes the ARG inference problem easier. From each marginal tree, we produce a list of tuples (*α, β, t*_*coal*_, *w*) where *α, β* are demes, *t*_*coal*_ is a particular coalescence time, and *w* is the weight of the coalescence (always 1 in our experiments). Let *T*_*c*_ be the list of all such values across all sampled trees and all processed chromosomes. Given *t*_*s*_, the number of time slices to use, we choose the corresponding time bin boundaries *τ*_*k*_ so that each time slice has roughly the same number of coalescences (∑*w*) from *T*_*c*_.

Bootstrapping trees is then performed by sampling with replacement from *T*_*c*_. Bootstrapping can be used for three purposes:

- The resulting 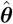 is derived from a coalescence count matrix **C** that is the average of the **C** matrices from the bootstrapped samples.
- When the **J** matrix is needed (for example when computing confidence intervals via the GIM-based method) we use the bootstrapped samples to estimate the expectation of the score function, as is typical.
- Bootstrapped samples can be used to compute confidence intervals directly, by solving the maximum likelihood problem for each sample and computing the sample standard deviation.

All our results are presented with *balanced* sample selection: all populations have the same number of samples.

### 4.4 Confidence Intervals

*mrpast* provides two methods for calculating parameter confidence intervals: computing the sample standard deviation from bootstrapped samples (where bootstrapping is over trees from the ARG(s)), or calculating the theoretical standard deviation via the Godambe Information Matrix (GIM), similar to [10].

The bootstrapping process involves running the full optimization pipeline (after creation of the coalescence matrices) for every sample. All results in this paper use 100 bootstrap samples. For models with few demes/parameters (such as the out-of-Africa models) bootstrapping can be reasonably fast (on the order of a few hours or less). For larger models, bootstrapping may be infeasible due to required compute time, in which case the GIM-based method can be used.

### 4.5 1,000 Genomes Data

We used the high-coverage 1,000 Genomes Data [8] and selected the first (based on VCF header order) 35 individuals from each of the populations YRI, CEU, CHB, CLM, PUR, and MXL. We then down-sampled the 1,000 Genomes Data to create four datasets: (YRI, CEU, CHB); (YRI, CEU, CHB, CLM); (YRI, CEU, CHB, PUR); (YRI, CEU, CHB, MXL). Only SNPs were retained, and non-segregating sites were removed. *tsinfer* performs polarization based on the ancestral sequence (GRCh38 ancestral sequence from Ensembl Release 112 [23]), unless there is no known ancestral state, in which case *tsinfer* imputes the state after local tree construction.

We used the data from bkgd [34] to avoid sampling ARG local trees in putatively non-neutral regions, in addition to the sampling described in section 4.3.2. We set a threshold of *B* ≥ 0.8, and only sample a tree if *B* ≥ 0.8 and the recombination rate ≤ 1 *×* 10^−9^ and the trees are at least 125Kbp apart.

Additionally, we confirmed experimentally that the effects of Neanderthal introgression (unaccounted for in the out-of-Africa models we have used) do not appear to affect our inference results. See Supplemental Section 1 for more details.

### 4.6 Time Discretization

The number of time slices can affect computation time (of the maximum-likelihood solver) and accuracy of the results. By default, *mrpast* automatically generates time slices, where the time boundaries are found by equally distributing all pairwise coalescence counts (ignoring population labels) among the number of time slices requested. In general, models with more epochs need more time slices, and the presence of the growth rate parameters requires more time slices for the growth approximation to be effective. In practice, some demographic models will have more parameters of interest in epochs close to the present time, and these epochs are covered by a lower proportion of the overall coalescence events. In these cases, the automatically generated time slices may be too coarse in this region to allow *mrpast* to accurately recover these recent-time parameters. Users may remedy this through specifying “left-skewed” time slices that are disproportionally more granular closer to present time, or through specifying additional timestamps (in generations) that divide the epoch(s) of interest. See Supplemental Table S1 for details of the time slices used for each result above.

Left-skewed time slices for *mrpast* are generated by continually bifurcating the time axis, choosing the left time slice for refinement each time. To compute *k* time slices, we start with time range (0, *τ*_*k*_), where *τ*_*k*_ = inf. Find time *τ*_*k*−1_ such that (0, *τ*_*k*−1_) and (*τ*_*k*−1_, *τ*_*k*_) have approximately equal number of panmictic coalescence events. Continue the same procedure with (0, *τ*_*k*−1_), finding the half-way point *τ*_*k*−2_, and so on until the algorithm terminates after finding *τ*_1_. The discretization of the time axis is then 0, *τ*_1_, *τ*_2_, …, *τ*_*k*_.

## Acknowledgement

The authors thank Rasmus Nielsen for helpful discussions during the early phase of this project, and Jinmin Li for helpful feedback on the manuscript. This work used Bridges-2 at Pittsburgh Supercomputing Center through allocation BIO230184 to X.W. from the Advanced Cyberinfrastructure Coordination Ecosystem: Services & Support (ACCESS) program, which is supported by National Science Foundation grants #2138259, #2138286, #2138307, #2137603, and #2138296. Z.J. was supported by the Nexus Scholars Program, College of Arts and Science, Cornell University. L.S. was supported by a JSPS KAKENHI grant [24K23946]. X.W. was supported by the National Institutes of Health grant [R35GM150579].

## Code Availability

The *mrpast* software is available at https://github.com/aprilweilab/mrpast.

