## Supplemental Text, Tables, Figures for "Inference of complex demographic history using composite likelihood based on whole-genome genealogies"

### Supplementary material: Joint parametric inference of complex demographic history using composite likelihood based on whole-genome genealogy

Drew DeHaas[1] Zhibai Jia[1] Leo Speidel[2] Xinzhu Wei[1*]

[1]Department of Computational Biology, Cornell University
[2]iTHEMS, RIKEN
*

**4 Tables** **6**

**5 Figures** **15**

#### 1 Neanderthal Admixture

The *OOA3G09* model, *OOA3G09_AM*, and American admixture models do not account for Neanderthal admixture, and we want to provide evidence that ignoring this event does not affect overall results. We modified the OutOfAfrica_2T12 [8] model to include a third deme representing Neanderthals (NEAN), in addition to AFR and EUR. Our modifications are based on the *OutOfAfricaExtendedNeandertalAdmixturePulse_3I21* model [2] parameters. We added a new epoch to *OutOfAfrica_2T12* to capture the population split between AFR and NEAN, and then added unidirectional migration from NEAN to EUR in the epoch spanning generations 920 - 2040 (approximately 30kya), see Figure S4. We used a migration rate of $2.67 \times 10^{-5}$, which is equal to taking the 3% total migration from the admixture model and dividing it equally across the epoch.

As seen in Figure S5, the *mrpast* inference using *OutOfAfrica_2T12* model is not negatively affected by the influence of this (ancient) Neanderthal introgression. When we try to infer the migration rate and time of this introgression, without using NEAN samples (i.e., NEAN is a ghost population in Fig. S4b), our results are more inaccurate (Fig. S5ac) than results ((Fig. S5bd)) from the simpler "misspecified" model (without the NEAN introgression modeled, Fig. S4a). Note that all of the results in Figure S5 use the genotype data generated with Neanderthal introgression (Fig. S4b), we are just varying whether the *inference* model tries to account for the introgression or not. These results imply that it is reasonable to use *mrpast* with out-of-Africa models to infer demographic parameters without explicitly accounting for Neanderthal introgression.

#### 2 Estimating Confidence Intervals

*mrpast* provides two methods for estimating parameter confidence intervals: computing the sample standard deviation from bootstrapped samples (where bootstrapping is over trees from the ARG(s)), or calculating the theoretical standard deviation via the Godambe Information Matrix (GIM), similar to [1].

The bootstrapping process involves running the full optimization pipeline (after creation of the coalescence matrices) for every sample. By default, we use 100 bootstrap samples. For models with few demes/parameters (such as the out-of-Africa models) bootstrapping can be reasonably fast (on the order of a few hours). For larger models, like the stepping stone model presented in the main text, bootstrapping is infeasible due to required compute time. A single optimization run (for a single replicate) for this large model times out after 24 hours (by default), so performing 100 bootstrap samples of 10 replicates each would be significant. The GIM-based confidence intervals are much cheaper, computationally, and especially useful for large models.

We found the GIM-based confidence intervals to be sensitive to numerical errors. In particular, we found our results to be better after implementing Ridder's method [6] for more accurate derivatives in the calculation of the $J$ and $H$ matrices.

The GIM $G$ is defined as $G = HJ^{-1}H$, where $H$ is the expectation (with respect to $\theta$) of the second-order partial derivatives of the likelihood function, and $J$ is the expectation of the first-order derivatives (see, for example, [5]). As is standard, we compute $H$ at $\hat{\theta}$ to get the expectation, and we use the average of $J$ over bootstrapped samples to get its expectation. From $G$ we can get the theoretical standard deviation of our likelihood function via $\sqrt{\operatorname{diag} G^{-1}}$. In practice, invertibility of these matrices can be challenging due to numerical issues, especially as the number of model parameters increases. Since for confidence intervals we only use $G^{-1}$, we can rearrange $G = HJ^{-1}H$ to be $G^{-1} = H^{-1}JH^{-1}$ so that only one matrix inverse operation is needed. We also achieved more consistent invertibility by rescaling the epoch times to be in units of "millions of generations", so that the scale is more similar to the other rate-based parameters.

#### 2.1 Population Growth Model: Time Slice Effects

In an attempt to improve the accuracy of the single population model with exponential growth, we added extra time slices at generations $50, 100, 150, 200, 250, 300$. This caused the simulated ARG results to be extremely accurate (all relative errors less than 5%, see Figure S9**b**, **c**). It also caused the variation to shrink drastically in both the replicated results for simulated ARGs and the bootstrapped results for inferred ARGs (Figure S9**d**, **e**). We speculate that adding these time slices causes the likelihood function to be more constrained to solutions that match the expected coalescence distribution at those short, very recent, time frames. For the simulated ARGs, the observed coalescences in those time slices match theory very well (hence the accuracy of inference), for inferred ARGs there may be bias or inaccuracy in those very small windows of time.

### 3 Likelihood Function

An ARG encapsulates the order of coalescence events among a set of haploid samples as well as the amount of time between each such event. A full likelihood function for an ARG, given a demographic model, is computationally intractable to work with. Instead, we assume independence between each observed pairwise coalescence and each sampled coalescent tree to formulate a composite likelihood function based on a subset of the information from the ARG. In practice, we utilize sparsely sampled trees along the genome that are at least 125 kilo-base-pairs (KBP) apart to make them relatively independent from each other.

We take an input model $G = \{\mathbf{M(1)}, \mathbf{\Lambda(1)}, \mathbf{T}_1, ..., \mathbf{M(E)}, \mathbf{\Lambda(E)}, \mathbf{T}_E\}$ representing the demographic structure being explored. Here, $E$ is the number of epochs. Given a model that has $D$ demes, each epoch $1 \leq e \leq E$ has a $D \times D$ migration rate matrix $\mathbf{M(e)}$, a $D$-length coalescent rate vector $\mathbf{\Lambda(e)}$, and an epoch end-time parameter $\mathbf{T}_e$. Note that $\mathbf{T}_E = \infty$ and is constant. The parameter vector $\boldsymbol{\theta}$ is a numeric vector that represents the unique non-zero parameters from $G$. If $\mathbf{M(e)}_{(\alpha,\beta)} = 0$, then there is no migration from demes $\alpha$ to $\beta$ in epoch $e$. If $\mathbf{M(e)}_{(\alpha,\beta)} = x$ ($x \in \mathbb{Z}$), then parameter $\boldsymbol{\theta}_x$ (the $x$-th element in $\boldsymbol{\theta}$) represents the migration rate from deme $\alpha$ to deme $\beta$ backward in time. $\mathbf{\Lambda(e)}_\alpha$ is defined similarly but represents coalescent rate of deme $\alpha$ in epoch $e$. Handling of growth rate parameters is discussed separately in the Supplemental Section 3.3.

Given a pair of lineages $i, j$, their locations in the $D$ demes can be specified by one of $\frac{D(D+1)}{2}$ states (all unordered combinations of deme pairs). We define a continuous-time Markov chain with $S = \frac{D(D+1)}{2} + 1$ states that represent both location and coalescent state of a pair of lineages. Each state

$\Gamma(\alpha,\beta) \in \{1, ..., \frac{D(D+1)}{2}\}$ represents one of the disjoint lineages $i$, $j$ being located in deme $\alpha$ and the other disjoint lineage being located in deme $\beta$. $\Gamma_{coal} = S = \frac{D(D+1)}{2} + 1$ is the absorbing state representing coalescence. Coalescence between lineages $i$, $j$ can only occur when two lineages are in the same deme (i.e., $\alpha = \beta$).

To enhance the resolution within each epoch, we further discretize time into $t = t_s + E$ bins, where $t_s$ is the number of additional discrete time slices. The ordered discrete times $\tau_1, ..., \tau_t$ are in units of generations backwards in time representing the end of each time bin, where $\tau_t = \infty$. Consequently, bin 1 is $[0, \tau_1)$ and bin $t$ is $[\tau_{t-1}, \infty)$. Each epoch, which may span multiple time bins, can have a different demographic topology and parameters, so we generate $E$ different Markov chains $MC(\mathbf{M(e)}, \mathbf{\Lambda(e)}, \mathbf{T}_e)$.

Consider a single epoch $e$. Given that a particular lineage pair $i, j$ is currently in state $\Gamma(\alpha,\beta)$ the possible transitions are $\Gamma(\alpha,\beta) \to \Gamma(\alpha,\beta)$, $\Gamma(\alpha,\beta) \to \Gamma(\gamma,\beta)$, $\Gamma(\alpha,\beta) \to \Gamma(\alpha,\gamma)$, and $\Gamma(\alpha,\beta) \to \Gamma_{coal}$. Transitions $\Gamma(\alpha,\beta) \to \Gamma(\gamma,\beta)$ and $\Gamma(\alpha,\beta) \to \Gamma(\alpha,\gamma)$ are defined for pairs $\alpha, \gamma$ ($\beta, \gamma$, respectively) only if the input model allows for migration between the corresponding demes (e.g., $\mathbf{M(e)}_{(\alpha,\gamma)} \neq 0$). From our $E$ input $\mathbf{M(e)}$ and $\mathbf{\Lambda(e)}$, we construct $E$ infinitesimal rate matrices ("$Q$-matrices" [4]) $\mathbf{Q(e)}$, of shape $S \times S$. The transition $\Gamma(\alpha,\beta) \to \Gamma(\alpha,\gamma)$ ($\alpha \neq \beta; \beta \neq \gamma$) has rate $\boldsymbol{\theta}_{\mathbf{M(e)}_{(\beta,\gamma)}}$. $\Gamma(\alpha,\alpha)$ can transition to the coalescence state $\Gamma_{coal}$ at rate $\boldsymbol{\theta}_{\mathbf{\Lambda(e)}_{\alpha}}$ or transition to $\Gamma(\alpha,\beta)$ ($\alpha \neq \beta$) at rate $2 \times \boldsymbol{\theta}_{\mathbf{M(e)}_{(\alpha,\beta)}}$ (because *either* lineage $i$ or $j$ can move to $\beta$). The diagonal element $\mathbf{Q(e)}_{(\Gamma(\alpha,\beta),\Gamma(\alpha,\beta))}$ is defined in the usual way as the negative sum of the nonzero off-diagonal elements of row $\mathbf{Q(e)}_{(\Gamma(\alpha,\beta),:)}$. The rate matrix $\mathbf{Q(e)}$ is used to compute state transition probabilities over time $\tau_k$ in epoch $e$ via $\mathbf{P_Q(k)} = e^{\mathbf{Q(e)}(\tau_k - \mathbf{T}_{e-1})}$. $\mathbf{P_Q(k)}$ has the following structure:

$$\mathbf{P_Q(k)} = \begin{Bmatrix} \mathbf{P}_\ell\mathbf{(k)} & \mathbf{P_c(k)} \\ \mathbf{0} & 1 \end{Bmatrix}$$

where $\mathbf{P}_\ell\mathbf{(k)}$ is a $(S-1) \times (S-1)$ matrix and $\mathbf{P_c(k)}$ is a $(S-1) \times 1$ matrix. $\mathbf{P}_\ell\mathbf{(k)}$ represents the probability of lineage $i, j$ being at a particular location at time $\tau_k$, *relative to the locations at the start of the current epoch*, and $\mathbf{P_c(k)}$ the cumulative probability of having coalesced between times $\mathbf{T}_{e-1}$ and $\tau_k$.

Let $\mathbf{L(e)}$ be the $(S-1) \times (S-1)$ matrix representing the location probabilities at the start of epoch $e$. $\mathbf{L(0)} = \mathbf{I}_{S-1}$, and $\mathbf{L(e)} = \prod_{i=0}^{e-1} \mathbf{L(i)}$, with the additional step that each product result must be normalized so the rows sum to 1. Only the latest $\mathbf{L(e)}$ is needed, so we keep a single $\mathbf{L}$ and update it if the current discrete time $\tau_k$ is the end of an epoch.

Let $\mathbf{\Omega}(e)$ be the vector of probabilities that a lineage pair starting in a particular state have coalesced in the epoch $e$. $\mathbf{\Omega}(0) = \mathbf{0}$. Similar to $\mathbf{L}$, we only need to compute a single $\mathbf{\Omega}$ for the previous epoch as we iterate the epochs in order. Since the current Markov chain only applies to the current epoch, $\mathbf{P_c(k)}$ needs to be adjusted by the probability that lineages have already coalesced in previous epochs, hence the need for $\mathbf{\Omega}$.

We calculate the cumulative coalescence probability matrix $\mathbf{F}$ row-by-row, where $\mathbf{F}_{(k,:)}$ is computed if $\tau_k$ is a time slice (see Supplemental Algorithm 1). Each discrete time $\tau_k$ can be a time slice and/or an epoch transition time (if $\tau_k = \mathbf{T}_e$ for any epoch $e$).

Given $\mathbf{F}$, the $t_s \times (S-1)$ matrix of cumulative coalescence probabilities, we convert to the probability density matrix $\mathbf{P}$, by setting $\mathbf{P}_{(t,:)} = \mathbf{1}$ and computing $\mathbf{P}_{(k,:)} = \mathbf{P}_{(k+1,:)} - \mathbf{F}_{(k,:)}$, for all $1 \leq k < t$.

Finally, we define the coalescence count matrix $\mathbf{C}$ to capture the observed coalescence counts between lineages in the ARG. Let each element $\mathbf{C(u)}_{(k,\Gamma(\alpha,\beta))} = \sum_{i\in\alpha,j\in\beta} I(\tau_{k-1} < T_{coal}(u,i,j) \leq \tau_k)$ where $i, j$ are samples, $i \in \alpha$ means $i$ is in deme $\alpha$ in the dataset, $u$ is a sampled tree from the ARG, and $I$ is the indicator function. Then $\mathbf{C} = \sum_u \mathbf{C(u)}$ over all sampled trees $u$. See Supplementary Section 3 for algorithms and more details, including admixture and growth rate handling.

*mrpast* uses the Subplex ([7]) solver from NLOpt ([3]) to solve for $\hat{\boldsymbol{\theta}}$ numerically. To find $\hat{\boldsymbol{\theta}} = argmin_{\boldsymbol{\theta}}(-cl(\boldsymbol{\theta}; y))$, we use multiple solver replicates with random initialization of $\boldsymbol{\theta}$ to ensure the global optimum. We use 10 solver replicates per epoch, e.g. a model with 2 epochs will use 20 solver replicates.

#### 3.1 Basic Algorithm

Algorithm 1 shows the basic algorithm for computing the coalesence probabilities for a model and parameter set. This simplified version is presented for clarity, see the following sections for more complex details.

#### 3.2 Population Splits and Admixture

**Algorithm 1** Theoretical coalescence probabilities

$\mathbf{L} \leftarrow \mathbf{I_{S-1}}$
$e \leftarrow 0$
$\boldsymbol{\Omega} \leftarrow \mathbf{0}$
**for** $k \leftarrow 1$ to $t$ **do**
  $\mathbf{P_Q(k)} \leftarrow e^{\mathbf{Q(e)} \times (\tau_k - T_{e-1})}$
  $\mathbf{P}_\ell(\mathbf{k}), \mathbf{P_c(k)} \leftarrow$ decompose($\mathbf{P_Q(k)}$)
  $\mathbf{P_c(k)} \leftarrow \mathbf{L} \times \mathbf{P_c(k)}$
  **if** isTimeslice($\tau_k$) **then**
    $\mathbf{F_{(k,:)}} \leftarrow ((\boldsymbol{\Omega} + \mathbf{P_c(k)}) - (\boldsymbol{\Omega} \times \mathbf{P_c(k)}))^T$
  **end if**
  **if** isEpoch($\tau_k$) **then**
    $\mathbf{L} \leftarrow$ rownorm($\mathbf{P}_\ell(\mathbf{k}) \times \mathbf{L}$)
    $\boldsymbol{\Omega} \leftarrow (\boldsymbol{\Omega} + \mathbf{P_c(k)}) - (\boldsymbol{\Omega} \times \mathbf{P_c(k)})$
    $e \leftarrow e + 1$
  **end if**
**end for**

A common modeling approach is to have an ancestral population that splits into multiple populations at a specific point in time. We require this time to be at an epoch boundary. We can achieve this by keeping a map $\overrightarrow{\Phi}$ from deme in the current epoch $e$ to the first epoch 0 (backwards in time). However, what follows is the more general approach of treating population splits as an admixture event with a proportion of 1.

Disallowing a deme to exist (backwards in time) after an admixture event simplifies the modeling. Specifically, we don't have to worry about circular relationships between a deme $A$ and $B$ where $A$ is an admixture of $B$ and $B$ is an admixture of $A$, because (a) all admixtures occur instantaneously at the start of the epoch and (b) demes terminate after admixture.

For each epoch, we take as input an admixture matrix $\mathbf{X}$, where at each cell $\mathbf{X}_{ij}$ we have either a constant value in range $[0, 1]$ or a symbol $\theta_i$ representing a parameter. Each row $i$ is the derived deme, and each column $j$ is the ancestral deme. So the value $0 \le p \le 1$ at $\mathbf{X}_{ij}$ is the proportion of deme $i$ that came from population $j$ (forward in time) at the given epoch.

A population split $i- > j$ (deme $i$ is derived from deme $j$) is just represented by $\mathbf{X}$ where the row $i$ has a single non-zero entry, and that entry is a value 1 in column $j$.

When $\mathbf{X} = \mathbf{I}_D$ (where $D$ is number of demes), there is no admixture (or population splits).

While the user specifies $\mathbf{X}$, we need to generate $\mathbf{S}$, which is the transformation of $\mathbf{X}$ from the deme-space to the Markov state-space (pairwise lineages). Properties of this transformation:

1. If $\mathbf{X}$ is the identity matrix, then $\mathbf{S}$ should be the identity matrix.
2. Consider a cell $\mathbf{S}_{\alpha\beta}$ where $\alpha = i, j$ and $\beta = k, l$. Then $\mathbf{S}_{\alpha,\beta}$ should be 0 unless *both* $\mathbf{X}_{ik}$ and $\mathbf{X}_{jl}$ (or both $\mathbf{X}_{ik}$ and $\mathbf{X}_{jk}$) are non-zero.
3. Each row is $\mathbf{S}$ should sum to 1.

For the mapping from $\mathbf{X}$ to $\mathbf{S}$, consider two states $\alpha = i, j$ and $\beta = k, l$. To compute $AS_{\alpha,\beta}$ we consider four cases:

1. $i = j$, $k = l$: $\mathbf{S}_{\alpha,\beta} = \mathbf{X}_{ik}^2$
2. $i = j$, $k \neq l$: $\mathbf{S}_{\alpha,\beta} = 2\mathbf{X}_{ik}\mathbf{X}_{il}$
3. $i \neq j$, $k = l$: $\mathbf{S}_{\alpha,\beta} = \mathbf{X}_{ik}\mathbf{X}_{jk}$
4. $i \neq j$, $k \neq l$: $\mathbf{S}_{\alpha,\beta} = \mathbf{X}_{ik}\mathbf{X}_{jl} + \mathbf{X}_{il}\mathbf{X}_{jk}$

We currently restrict each row (derived deme) of $\mathbf{X}$ to be made up entirely of symbolic proportions or constant proportions. In both cases the proportions in the row have to sum to 1, so in the symbolic case where there are $k$ parameters representing proportions there are only $k - 1$ degrees of freedom. The numerical optimization only uses the first (arbitrary order) $k - 1$ parameters.

Algorithm 2 shows how we apply the $\mathbf{S}(e)$ matrix for each epoch. Since we process the epochs in order, we keep a cumulative admixture matrix $\mathbf{A}$ that represents the application of $\mathbf{S}(0) \times \mathbf{S}(1) \times ... \times \mathbf{S}(e)$ where $e$ is the current epoch and $\mathbf{S}(0)$ is the identity.

**Algorithm 2** Theoretical coalescence probabilities: with population split admixture

$\mathbf{A} \leftarrow \mathbf{I_{S-1}}$
$\mathbf{L} \leftarrow \mathbf{I_{S-1}}$
$e \leftarrow 0$
$\mathbf{\Omega} \leftarrow \mathbf{0}$
**for** $k \leftarrow 1$ to $t$ **do**
  $\mathbf{P_Q(k)} \leftarrow e^{\mathbf{Q(e)} \times (\tau_k - T_{e-1}))}$
  $\mathbf{P}_\ell(\mathbf{k}), \mathbf{P_c(k)} \leftarrow \text{decompose}(\mathbf{P_Q(k)})$
  $\mathbf{P_c(k)} \leftarrow \mathbf{L} \times \mathbf{P_c(k)}$
  **if** isTimeslice($\tau_k$) **then**
    $\mathbf{F_{(k,:)}} \leftarrow ((\mathbf{\Omega} + \mathbf{P_c(k)}) - (\mathbf{\Omega} \times \mathbf{P_c(k)}))^T$
  **end if**
  **if** isEpoch($\tau_k$) **then**
    $\mathbf{L} \leftarrow \text{rownorm}(\mathbf{L} \times \mathbf{A} \times \mathbf{P}_\ell(\mathbf{k}))$
    $\mathbf{A} \leftarrow \mathbf{A} \times \mathbf{S(e+1)}$
    $\mathbf{L} \leftarrow \mathbf{L} \times \mathbf{A}$
    $\mathbf{\Omega} \leftarrow (\mathbf{\Omega} + \mathbf{P_c(k)}) - (\mathbf{\Omega} \times \mathbf{P_c(k)})$
    $e \leftarrow e + 1$
  **end if**
**end for**

##### 3.3 Growth Rates

Within each epoch, we have a per-deme coalescence rate of $\lambda_i$ (lineage pairs/generation). At each time slice $\tau_k$ we solve the matrix exponential $e^{Q\tau_k}$ where $Q$ is the infinitesimal rate matrix encoding state transition for pairs of lineages. The state for when a pair of lineages are in the same deme has a transition to the coalescence state with a rate of $\lambda_i$ for deme $i$.

In order to incorporate growth rate, we introduce new growth parameters $\alpha_i$. Given diploid samples, we have $Ne_i^{orig} = \frac{1}{2\lambda_i}$. With growth rate $\alpha_i$ at time $\tau_k$ we have $Ne_i^{new} = Ne_i^{orig} \times e^{-\alpha_i \tau_k} = \frac{1}{2\lambda_i} \times e^{-\alpha_i \tau_k}$, which gives us updated coalescence rate $\lambda_i^{new} = \frac{1}{2Ne_i^{new}} = \lambda_i e^{\alpha_i \tau_k}$.

Each solution of $e^{Q\tau_k}$ is the cumulative probability from time 0 (the start of the current epoch) to $\tau_k$. To approximate the adjusted coalescence rate with our discretized time intervals, we use the average rate from 0 to $\tau_k$. We can take the integral $\int_0^{\tau_k} e^{\alpha_i \tau_k} = \frac{e^{\alpha_i \tau_k} - 1}{\alpha_i}$ and then average by dividing by $\tau_k$. At each $\tau_k$ then our coalescence rate (and thus transition rate to the coalescence state in the $Q$ matrix) for deme $i$ is $\lambda_i \frac{e^{\alpha_i \tau_k} - 1}{\alpha_i \tau_k}$

### 4 Tables

Table S1: Time slices used by models

| Model | Automatic slices | Left-skewed? | Manually specified slices |
|---|---|---|---|
| Stepping stone 4x5 | 20 | No | N/A |
| African $N_e$ | 200 | No | N/A |
| OOA3G09 | 50 | Yes | N/A |
| OOA3_AM | 200 | No | N/A |
| AA5 (Admixture) | 200 | No | [5, 10, 15, 20] |

The manually specified time slices column lists the times (in generations) for additional time slices that were added.

Table S2: *mrpast* inferred parameters for the OOA3 (symmetric migration) model.

| **Description** | **Epoch** | **Median Value** | **Standard Deviation** |
|---|---|---|---|
| Migration rate: YRI and CEU | 1 | $2.64 \times 10^{-4}$ | $3.21 \times 10^{-6}$ |
| Migration rate: YRI and CEU | 0 | $4.04 \times 10^{-5}$ | $1.15 \times 10^{-6}$ |
| Migration rate: YRI and CHB | 0 | $\mathbf{1.00 \times 10^{-5}}$ | $2.42 \times 10^{-17}$ |
| Migration rate: CEU and CHB | 0 | $1.20 \times 10^{-4}$ | $2.19 \times 10^{-6}$ |
| $N_e$ for YRI | 3 | 16073.02 | 198.92 |
| $N_e$ for YRI | 0 - 2 | 20695.69 | 451.34 |
| $N_e$ for CEU | 1 | 2805.06 | 17.62 |
| $N_e$ for CEU | 0 | 35179.32 | 388.32 |
| $N_e$ for CHB | 0 | 47136.84 | 599.54 |
| Growth rate for deme CEU | 0 | $2.62 \times 10^{-3}$ | $2.49 \times 10^{-5}$ |
| Growth rate for deme CHB | 0 | $3.46 \times 10^{-3}$ | $2.43 \times 10^{-5}$ |

Fixed epoch times were used. ARGs were inferred for CEU, CHB, and YRI samples in the 1000 Genomes Project. Summary of 100 bootstrap iterations. Bold values are on the lower or upper bound.

Table S3: *mrpast* inferred parameters for the OOA3 (asymmetric migration) model.

| **Description** | **Epoch** | **Median Value** | **Standard Deviation** |
|---|---|---|---|
| Migration rate: YRI to CEU | 1 | $1.63 \times 10^{-4}$ | $3.38 \times 10^{-6}$ |
| Migration rate: YRI to CEU | 0 | $2.48 \times 10^{-5}$ | $2.55 \times 10^{-6}$ |
| Migration rate: YRI to CHB | 0 | $3.70 \times 10^{-5}$ | $1.42 \times 10^{-6}$ |
| Migration rate: CHB to CEU | 0 | $\mathbf{1.00 \times 10^{-5}}$ | $1.05 \times 10^{-15}$ |
| Migration rate: CEU to YRI | 0 | $9.83 \times 10^{-5}$ | $4.46 \times 10^{-6}$ |
| Migration rate: CHB to YRI | 0 | $\mathbf{1.00 \times 10^{-5}}$ | $9.26 \times 10^{-18}$ |
| Migration rate: CEU to CHB | 0 | $1.94 \times 10^{-4}$ | $3.90 \times 10^{-6}$ |
| Migration rate: CEU to YRI | 1 | $3.27 \times 10^{-4}$ | $5.44 \times 10^{-6}$ |
| $N_e$ for YRI | 3 | 17311.43 | 225.67 |
| $N_e$ for YRI | 0 - 2 | 14571.52 | 139.90 |
| $N_e$ for CEU | 1 | 2200.57 | 25.17 |
| $N_e$ for CEU | 0 | 33218.15 | 345.93 |
| $N_e$ for CHB | 0 | 45729.58 | 584.60 |
| Growth rate for deme CEU | 0 | $2.65 \times 10^{-3}$ | $2.43 \times 10^{-5}$ |
| Growth rate for deme CHB | 0 | $3.20 \times 10^{-3}$ | $2.33 \times 10^{-5}$ |

Fixed epoch times were used. ARGs were inferred for CEU, CHB, and YRI samples in the 1000 Genomes Project. Summary of 100 bootstrap iterations. Migrations are described backwards in time. Bold values are on the lower or upper bound.

Table S4: *mrpast* inferred parameters for the OOA3 (asymmetric migration) model using FIN instead of CEU.

| Description | Epoch | Median Value | Standard Deviation |
|---|---|---|---|
| Migration rate: YRI to FIN | 1 | $2.03 \times 10^{-4}$ | $4.26 \times 10^{-6}$ |
| Migration rate: YRI to FIN | 0 | $6.89 \times 10^{-6}$ | $2.76 \times 10^{-6}$ |
| Migration rate: YRI to CHB | 0 | $5.25 \times 10^{-5}$ | $1.54 \times 10^{-6}$ |
| Migration rate: CHB to FIN | 0 | $\mathbf{1.00 \times 10^{-6}}$ | $5.15 \times 10^{-16}$ |
| Migration rate: FIN to YRI | 0 | $1.31 \times 10^{-4}$ | $6.30 \times 10^{-6}$ |
| Migration rate: CHB to YRI | 0 | $\mathbf{1.00 \times 10^{-6}}$ | $1.34 \times 10^{-16}$ |
| Migration rate: FIN to CHB | 0 | $5.12 \times 10^{-4}$ | $5.89 \times 10^{-6}$ |
| Migration rate: FIN to YRI | 1 | $3.35 \times 10^{-4}$ | $5.31 \times 10^{-6}$ |
| $N_e$ for YRI | 3 | 12950.82 | 114.40 |
| $N_e$ for YRI | 0 - 2 | 19782.14 | 262.98 |
| $N_e$ for FIN | 1 | 2163.55 | 24.12 |
| $N_e$ for FIN | 0 | 14995.03 | 128.57 |
| $N_e$ for CHB | 0 | 45610.80 | 577.19 |
| Growth rate for FIN | 0 | $1.89 \times 10^{-3}$ | $1.97 \times 10^{-5}$ |
| Growth rate for CHB | 0 | $3.24 \times 10^{-3}$ | $2.40 \times 10^{-5}$ |

Fixed epoch times were used. ARGs were inferred for FIN, CHB, and YRI samples in the 1000 Genomes Project. 200 time slices (not skewed). Summary of 100 bootstrap iterations. Migrations are described backwards in time. Bold values are on the lower or upper bound.

Table S5: *mrpast* inferred parameters for the OOA3 (asymmetric migration) model with an extra epoch at 0-100 generations

| **Description** | **Epoch** | **Median Value** | **Standard Deviation** |
|---|---|---|---|
| Migration rate: YRI to CEU | 2 | $1.65 \times 10^{-4}$ | $3.37 \times 10^{-6}$ |
| Migration rate: YRI to CEU | 0, 1 | $2.50 \times 10^{-5}$ | $2.42 \times 10^{-6}$ |
| Migration rate: YRI to CHB | 0, 1 | $3.79 \times 10^{-5}$ | $1.36 \times 10^{-6}$ |
| Migration rate: CHB to CEU | 1 | $\mathbf{1.00 \times 10^{-6}}$ | $4.92 \times 10^{-5}$ |
| Migration rate: CEU to YRI | 0, 1 | $8.91 \times 10^{-5}$ | $4.52 \times 10^{-6}$ |
| Migration rate: CHB to YRI | 0, 1 | $\mathbf{1.00 \times 10^{-6}}$ | $1.05 \times 10^{-17}$ |
| Migration rate: CEU to CHB | 1 | $2.20 \times 10^{-4}$ | $4.52 \times 10^{-5}$ |
| Migration rate: CEU to YRI | 2 | $3.27 \times 10^{-4}$ | $5.47 \times 10^{-6}$ |
| Migration rate: CHB to CEU | 0 | $\mathbf{1.00 \times 10^{-6}}$ | $2.21 \times 10^{-15}$ |
| Migration rate: CEU to CHB | 0 | $7.25 \times 10^{-5}$ | $8.15 \times 10^{-6}$ |
| $N_e$ for YRI | 4 | 17321.92 | 225.88 |
| $N_e$ for YRI | 0 - 3 | 14542.58 | 141.46 |
| $N_e$ for CEU | 2 | 2232.09 | 26.20 |
| $N_e$ for CEU | 0, 1 | 26302.64 | 232.95 |
| $N_e$ for CHB | 0, 1 | 33576.51 | 368.88 |
| Growth rate for CEU | 1 | $2.72 \times 10^{-3}$ | $7.62 \times 10^{-5}$ |
| Growth rate for CHB | 1 | $3.18 \times 10^{-3}$ | $7.51 \times 10^{-5}$ |

Fixed epoch times were used. ARGs were inferred for CEU, CHB, and YRI samples in the 1000 Genomes Project. 50 time slices (left skewed). Summary of 100 bootstrap iterations. Migrations are described backwards in time. Bold values are on the lower or upper bound. Epochs are (0-100), (100-848), (848-5600), (5600-8800), (8800-inf)

Table S6: *mrpast* inferred parameters for the AA5 (American admixture) model, using CLM.

| **Description** | **Epoch** | **Median Value** | **Standard Deviation** |
|---|---|---|---|
| End of epoch 0 | | 19.35 | 0.44 |
| End of epoch 1 | | 763.71 | 2.14 |
| End of epoch 2 | | 994.95 | 16.49 |
| End of epoch 3 | | 5467.91 | 33.59 |
| End of epoch 4 | | **9000.00** | $4.65 \times 10^{-12}$ |
| Migration rate: AFR and EUR | 3 | $2.97 \times 10^{-4}$ | $5.70 \times 10^{-6}$ |
| Migration rate: AFR and EUR | 0 - 2 | $5.32 \times 10^{-5}$ | $1.66 \times 10^{-6}$ |
| Migration rate: AFR and ASIA | 0 - 2 | $2.11 \times 10^{-5}$ | $1.26 \times 10^{-6}$ |
| Migration rate: EUR and ASIA | 0 - 2 | $1.98 \times 10^{-4}$ | $2.91 \times 10^{-6}$ |
| $N_e$ for AFR | 5 | 12692.48 | 97.20 |
| $N_e$ for AFR | 0 - 4 | 20185.22 | 288.02 |
| $N_e$ for EUR | 3 | 2868.72 | 26.35 |
| $N_e$ for EUR | 2 | 3141.68 | 54.33 |
| $N_e$ for ASIA | 2 | 1854.74 | 66.28 |
| $N_e$ for EUR | 0, 1 | 32563.69 | 611.01 |
| $N_e$ for ASIA | 0, 1 | 29617.77 | 699.10 |
| $N_e$ for NAT | 0, 1 | 1028.97 | 34.22 |
| $N_e$ for ADMIX | 0 | **100000.00** | $3.36 \times 10^{-5}$ |
| Growth rate for EUR | 1 | $2.16 \times 10^{-3}$ | $4.91 \times 10^{-5}$ |
| Growth rate for ASIA | 1 | $2.65 \times 10^{-3}$ | $6.13 \times 10^{-5}$ |
| Admixture: AFR and ADMIX | | **0.02** | $3.53 \times 10^{-3}$ |
| Admixture: EUR and ADMIX | | 0.80 | $5.27 \times 10^{-3}$ |
| Admixture: NAT and ADMIX | | 0.18 | $3.27 \times 10^{-3}$ |

ARGs were inferred for CLM, CEU, CHB, and YRI samples in the 1000 Genomes Project. Summary of 100 bootstrap iterations. Migrations are described backwards in time. Bold values are on the lower or upper bound.

Table S7: *mrpast* inferred parameters for the AA5 (American admixture) model, using PUR.

| **Description** | **Epoch** | **Median Value** | **Standard Deviation** |
|---|---|---|---|
| End of epoch 0 | | 19.77 | 0.48 |
| End of epoch 1 | | 791.47 | 22.46 |
| End of epoch 2 | | **1000.00** | 22.30 |
| End of epoch 3 | | 5594.01 | 56.06 |
| End of epoch 4 | | **9000.00** | 0.00 |
| Migration rate: AFR and EUR | 3 | $2.93 \times 10^{-4}$ | $5.84 \times 10^{-6}$ |
| Migration rate: AFR and EUR | 0 - 2 | $5.46 \times 10^{-5}$ | $1.92 \times 10^{-6}$ |
| Migration rate: AFR and ASIA | 0 - 2 | $2.29 \times 10^{-5}$ | $1.56 \times 10^{-6}$ |
| Migration rate: EUR and ASIA | 0 - 2 | $1.92 \times 10^{-4}$ | $8.79 \times 10^{-6}$ |
| $N_e$ for AFR | 5 | 12541.80 | 83.52 |
| $N_e$ for AFR | 0 - 4 | 20840.36 | 319.40 |
| $N_e$ for EUR | 3 | 2893.81 | 25.92 |
| $N_e$ for EUR | 2 | 3105.71 | 277.56 |
| $N_e$ for ASIA | 2 | 1715.86 | 175.82 |
| $N_e$ for EUR | 0, 1 | 35882.27 | 6440.94 |
| $N_e$ for ASIA | 0, 1 | 33398.58 | 6701.18 |
| $N_e$ for NAT | 0, 1 | 1250.97 | 104.69 |
| $N_e$ for ADMIX | 0 | 88928.41 | 2565.44 |
| Growth rate for EUR | 1 | $2.40 \times 10^{-3}$ | $2.50 \times 10^{-4}$ |
| Growth rate for ASIA | 1 | $2.88 \times 10^{-3}$ | $2.55 \times 10^{-4}$ |
| Admixture proportion: AFR and ADMIX | | 0.10 | $4.83 \times 10^{-3}$ |
| Admixture proportion: EUR and ADMIX | | 0.77 | 0.01 |
| Admixture proportion: NAT and ADMIX | | 0.13 | $7.40 \times 10^{-3}$ |

ARGs were inferred for PUR, CEU, CHB, and YRI samples in the 1000 Genomes Project. Summary of 100 bootstrap iterations. Migrations are described backwards in time. Bold values are on the lower or upper bound.

Table S8: *mrpast* inferred parameters for the AA5 (American admixture) model, using MXL.

| **Description** | **Epoch** | **Median Value** | **Standard Deviation** |
|---|---|---|---|
| End of epoch 0 | | 19.97 | $5.45 \times 10^{-3}$ |
| End of epoch 1 | | 733.62 | 1.24 |
| End of epoch 2 | | 978.64 | 2.02 |
| End of epoch 3 | | **5400.00** | $1.11 \times 10^{-12}$ |
| End of epoch 4 | | **9000.00** | $3.73 \times 10^{-12}$ |
| Migration rate: AFR and EUR | 3 | $2.91 \times 10^{-4}$ | $3.68 \times 10^{-6}$ |
| Migration rate: AFR and EUR | 0 - 2 | $5.35 \times 10^{-5}$ | $1.43 \times 10^{-6}$ |
| Migration rate: AFR and ASIA | 0 - 2 | $1.93 \times 10^{-5}$ | $9.09 \times 10^{-7}$ |
| Migration rate: EUR and ASIA | 0 - 2 | $2.21 \times 10^{-4}$ | $2.54 \times 10^{-6}$ |
| $N_e$ for AFR | 5 | 12787.59 | 114.84 |
| $N_e$ for AFR | 0 - 4 | 19214.62 | 255.38 |
| $N_e$ for EUR | 3 | 2839.82 | 21.18 |
| $N_e$ for EUR | 2 | 3084.83 | 43.27 |
| $N_e$ for ASIA | 2 | 2057.54 | 31.56 |
| $N_e$ for EUR | 0, 1 | 29825.06 | 429.34 |
| $N_e$ for ASIA | 0, 1 | 24857.62 | 401.58 |
| $N_e$ for NAT | 0, 1 | 1006.38 | 24.18 |
| $N_e$ for ADMIX | 0 | **100000.00** | $4.01 \times 10^{-6}$ |
| Growth rate for EUR | 1 | $1.97 \times 10^{-3}$ | $4.11 \times 10^{-5}$ |
| Growth rate for ASIA | 1 | $2.28 \times 10^{-3}$ | $4.25 \times 10^{-5}$ |
| Admixture proportion: AFR and ADMIX | | **0.02** | $3.49 \times 10^{-18}$ |
| Admixture proportion: EUR and ADMIX | | 0.69 | $2.87 \times 10^{-3}$ |
| Admixture proportion: NAT and ADMIX | | 0.29 | $2.87 \times 10^{-3}$ |

ARGs were inferred for MXL, CEU, CHB, and YRI samples in the 1000 Genomes Project. Summary of 100 bootstrap iterations. Migrations are described backwards in time. Bold values are on the lower or upper bound.

Table S9: Number of marginal trees in ARGs and the number of trees that *mrpast* sampled.

| **Model** | **Sim. Total** | **Sim. Sampled** | **Infer. Total** | **Infer. Sampled** |
|---|---|---|---|---|
| Stepping stone 4x5 | 15449063 | 10829 | N/A | N/A |
| African $N_e$ | 6677990 | 9745 | 1866029 | 11563 |
| OOA3G09 | 7524279 | 9951 | 2935046 | 11633 |
| AA5 (Admixture) | 8703483 | 10136 | 3377337 | 11638 |

For simulated and inferred ARGs for each model from simulated data (20 chromosomes each).

### 5 Figures

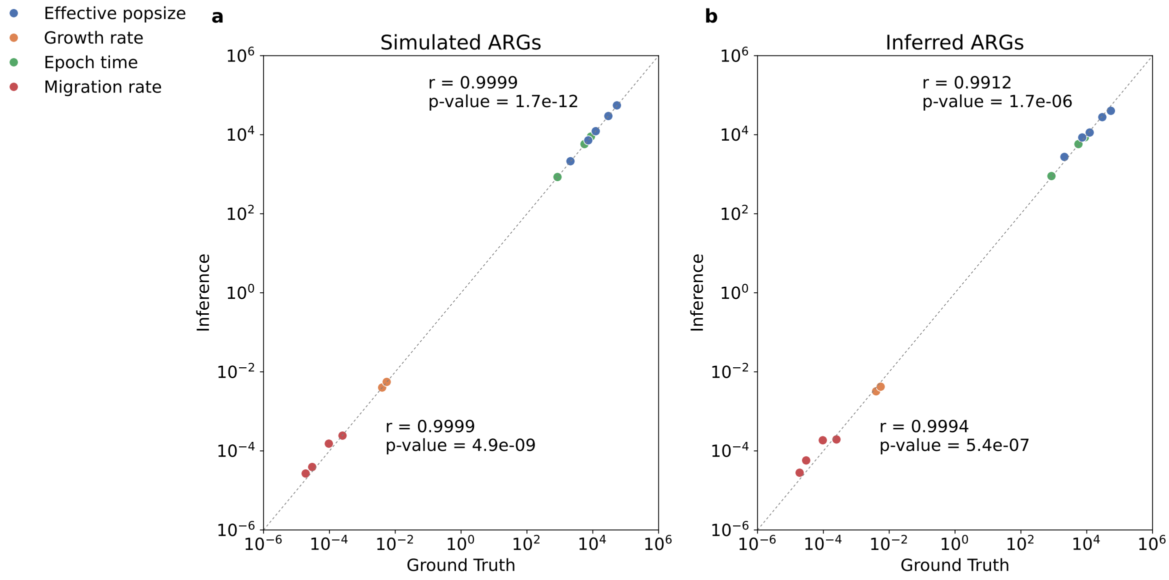

Figure S1: **OOA3G09 model, simulated data, 10 individuals per deme.** **a**: (Simulated ARGs) show the true (simulated) values against the inferred values on a log-log scale. The Pearson's $r$ and associated p-value are shown separately for the time and population size parameter (upper) and the rate parameters (lower). **b**: Same as **a**, but using inferred ARGs on simulated data.

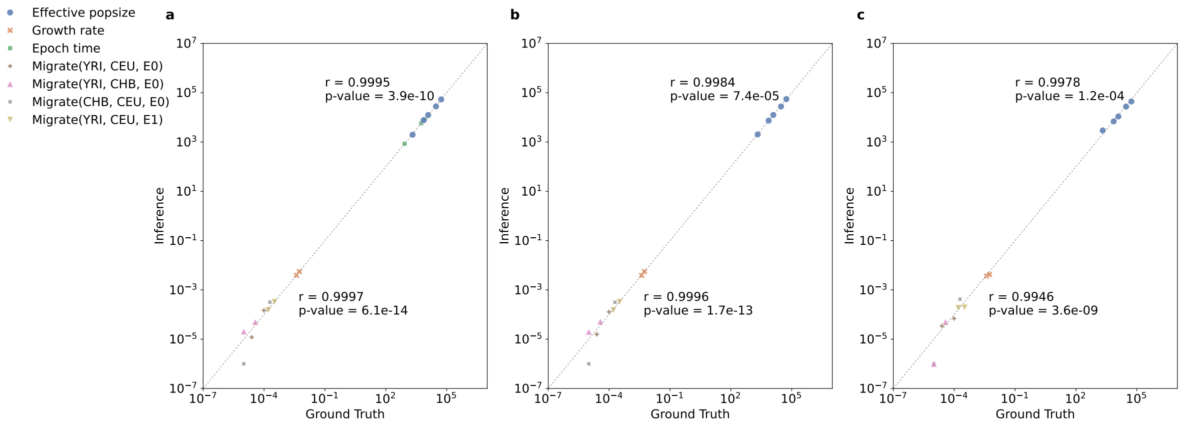

Figure S2: **OOA3G09 asymmetric model, simulated data.** **a**: (Simulated ARGs) show the true (simulated) values against the inferred values on a log-log scale. The epoch time parameters are free variables (found by the solver and plotted). The Pearson's $r$ and associated p-value are shown separately for the time and population size parameter (upper) and the rate parameters (lower). **b**: Same as **a**, except the epoch time parameters are fixed (constant values). **c**: Same as **b**, but using inferred ARGs on simulated data. All three results use 50 individuals per deme.

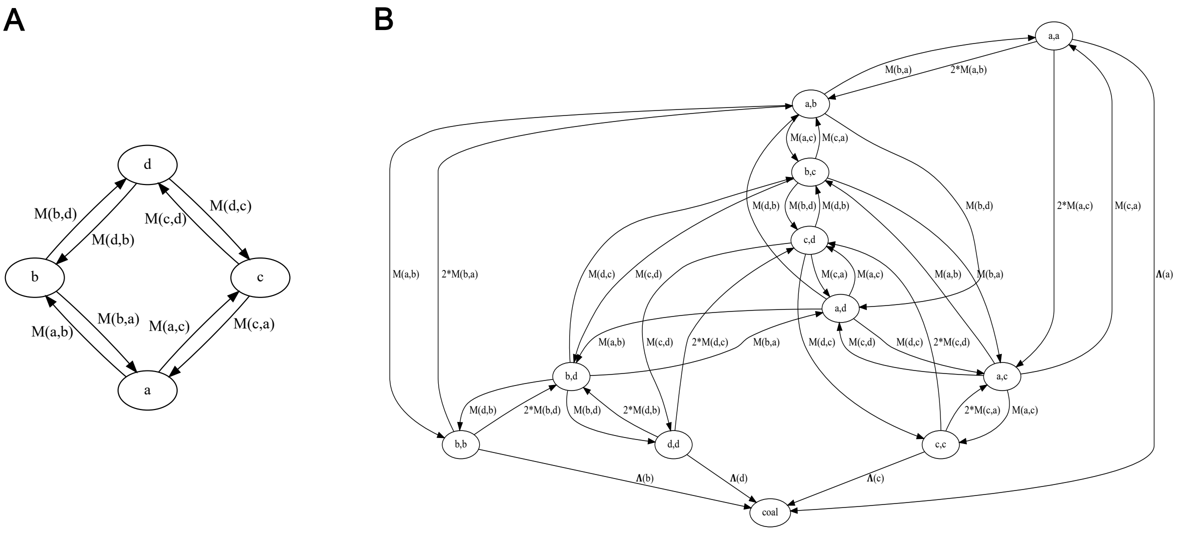

Figure S3: **Mapping from deme migration graph to Markov chain transition graph.** **A**: a simple 4-deme topology parameterized by 8 asymmetric migration rates. **B**: The corresponding Markov chain transitions, where each node is a state representing the current location of a lineage pair, and a single coalescence state that is only reachable when both lineages are in the same deme.

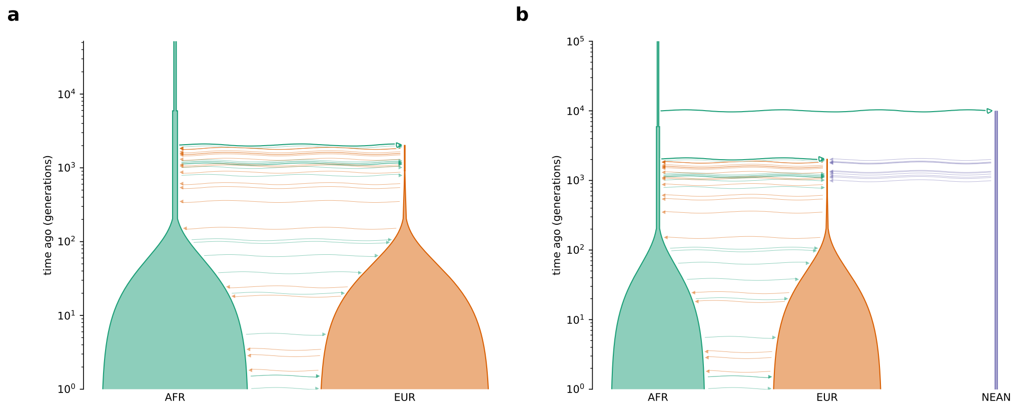

Figure S4: **Out-of-Africa models for testing the effect of neanderthal introgression on *mr-past* inference.** **a**: the OutOfAfrica_2T12 without neanderthal introgression (as used elsewhere in our testing). **b**: The same model, modified to have an additional epoch (to model the split of NEAN from AFR) and a continuous migration from NEAN to EUR that sums to 3% of the lineages.

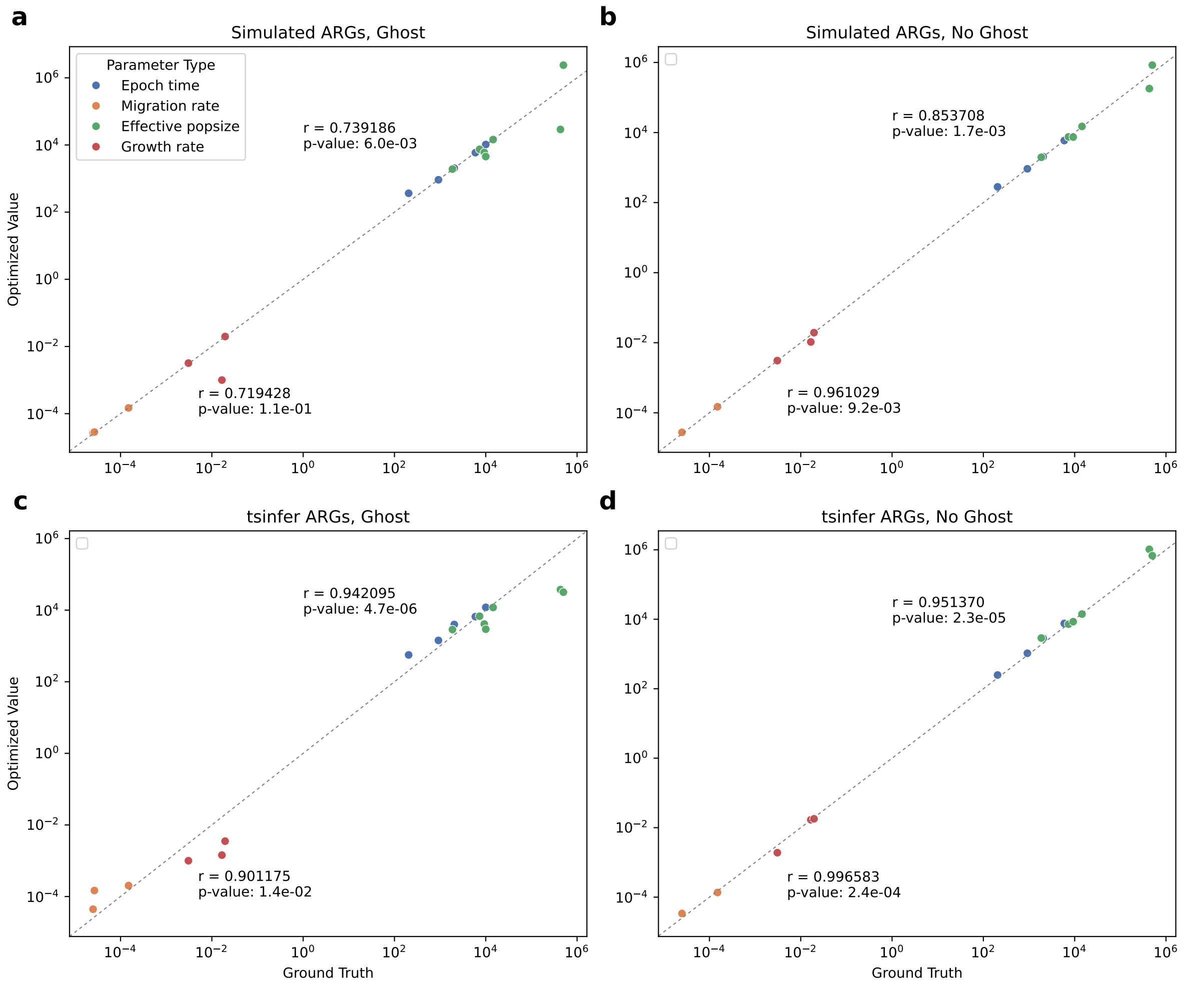

Figure S5: **Testing the effect of neanderthal introgression on *mrpast* inference.** All panels use data simulated with neanderthal introgression, and show *mrpast* inferred parameters plotted against their ground-truth values. **a**: Results for simulated ARGs containing only AFR and EUR samples, but with a model that attempts to infer parameters for the NEAN population (i.e., NEAN is a ghost population). **b**: Results for simulated ARGs containing only AFR and EUR samples, with a model that is only inferring AFR and EUR samples. **c**: Same as **a**, but with ARGs from tsinfer. **d**: Same as **b**, but with ARGs from tsinfer.

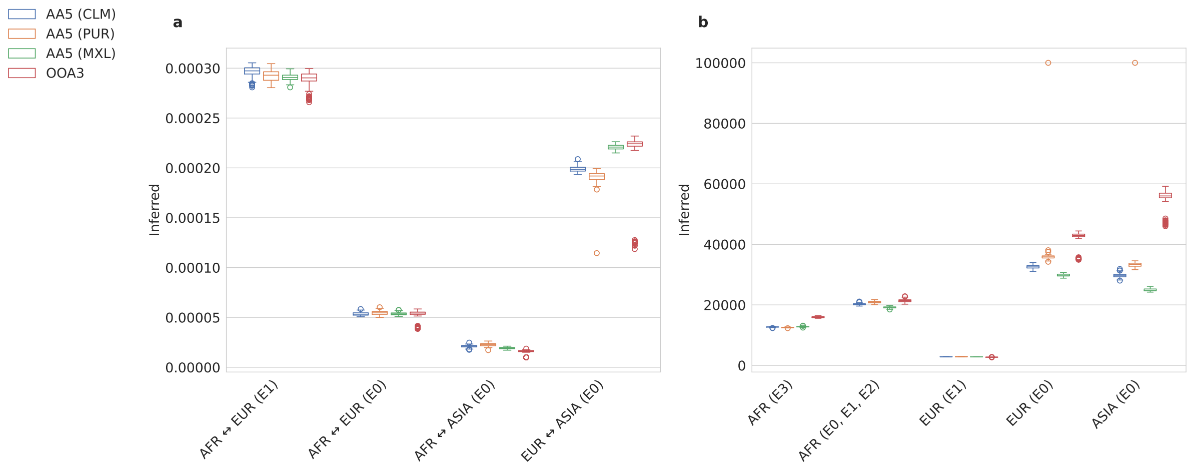

Figure S6: **Common parameters inferred by two different (nested) models.** The AA5 model extends the OOA3 model with additional parameters. Here we compare some of the common parameters: the migration rates (**a**) and effective population sizes (**b**). The epochs are specified as E0, E1, E2, and correspond to the OOA3 model epochs, not the AA5 epochs (of which there are more, to get finer granularity results).

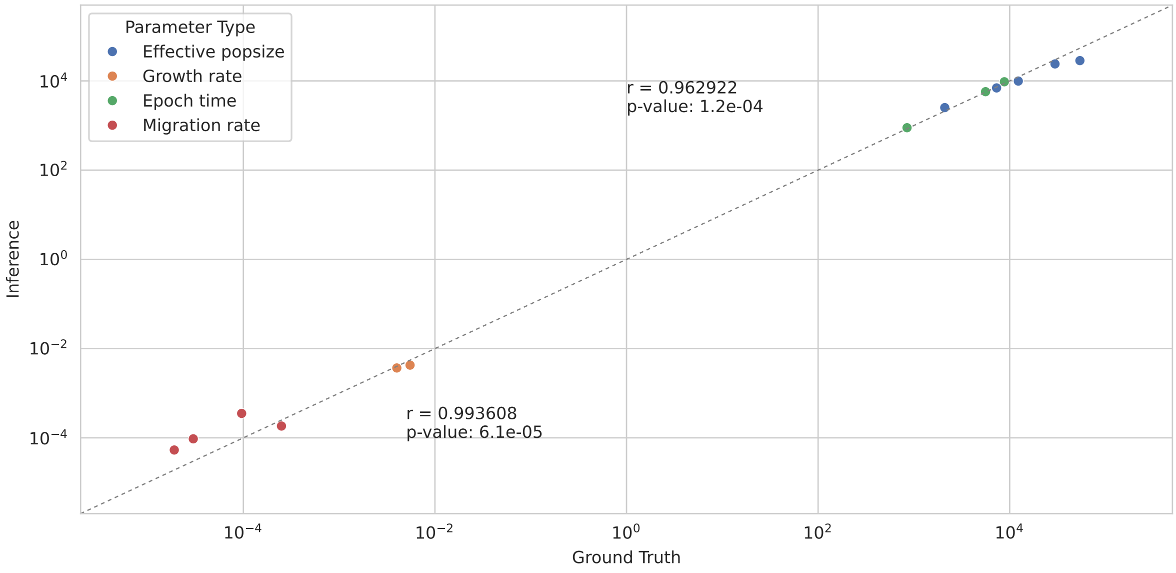

Figure S7: **Results using Relate-inferred ARGs.** Simulated data from the OOA3G09 model, inferred with Relate using 10 MC/MC samples for branch lengths.

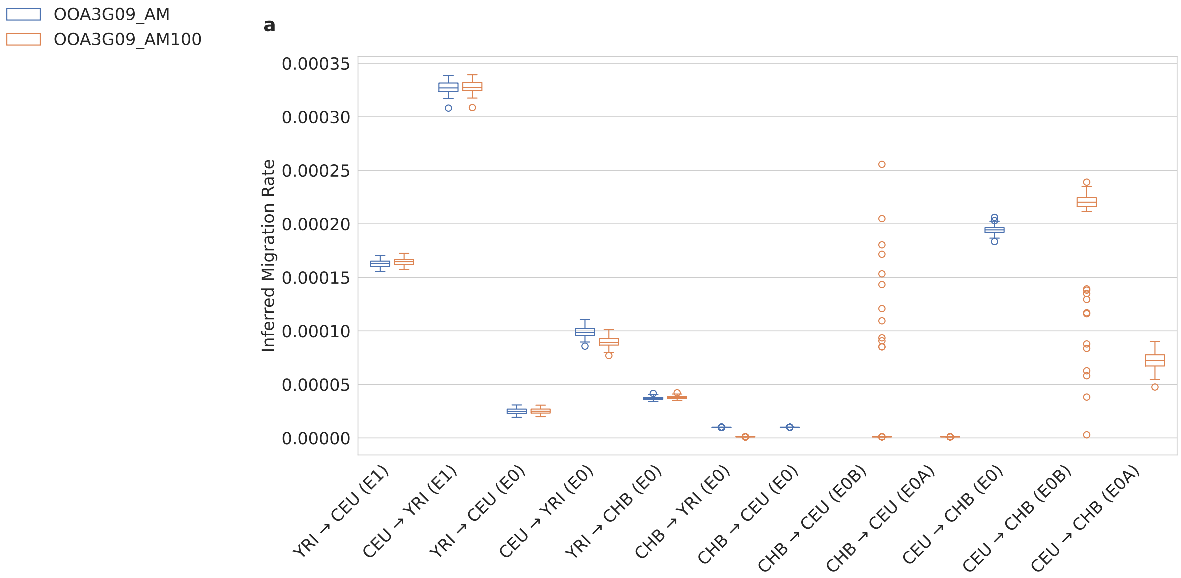

Figure S8: **Effect of adding another epoch to OOA3G09_AM.** The epochs (in generations) are E0 (0-848), E1 (848-5600), E2 (5600-8800), and E3 (8800-inf). OOA3G09_AM100 is the same model as OOA3G09_AM, except we split E0 into two parts, E0A (0-100) and E0B (100-848) and split the migration rate parameters involving CHB and CEU as well. This figure compares migration rate estimated on 1,000 Genomes Data between these two model variations.

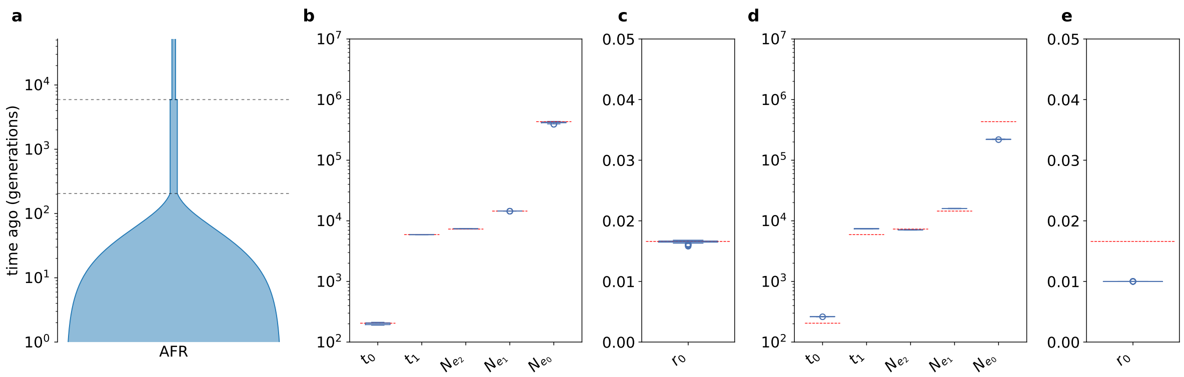

Figure S9: **Adding extra time slices to the population growth model.** The single population growth model with time slices added at times 50, 100, 150, 200, 250, 300. Diagram of the model in panel **a**, simulated ARG results in panels **b**, **c**, and inferred ARG results in panels **d**, **e**. The simulated ARGs show variation from 50 replicated simulations. The inferred ARGs show variation from 100 bootstrap samples.
